# Microtubule Lattice Spacing Governs MAP-Motor Regulation

**DOI:** 10.64898/2026.08.03.742653

**Authors:** Jonathan Fernandes, Joseph Slivka, Aryan Taheri, Qikai Zhang, Viola Zhao, Chuzida Chen, Parnika Kant, Ahmet Yildiz

**Affiliations:** Department of Chemistry, University of California at Berkeley, Berkeley, CA 94720 USA; Department of Physics, University of California at Berkeley, Berkeley, CA 94720 USA; Department of Molecular and Cell Biology, University of California at Berkeley, Berkeley, CA 94720 USA; Biophysics Graduate Group, University of California at Berkeley, Berkeley, CA 94720 USA

## Abstract

Microtubules (MTs) serve as intracellular tracks that enable molecular motors to transport cargos to specific cellular destinations. It has been proposed that the signals directing motor-driven transport are encoded on MTs through different isotypes, lattice conformations, and post-translational modifications (PTMs) of tubulin, or MT-associated proteins (MAPs) that decorate the MT surface. However, molecular predictions of these models have not been rigorously tested in vitro. Using isotypically pure recombinant tubulin and biochemical reconstitution, we examined how tubulin PTMs and MT lattice spacing influence MAP binding and kinesin-1 motility. We found that kinesin-1 is largely insensitive to tubulin PTMs but is strongly regulated by MT lattice spacing. Likewise, the MAPs tau, MAP7, MAP4, DCX, and MAP9 exhibit little sensitivity to tubulin PTMs, whereas the MT-binding affinities of tau, DCX, and MAP7 depend on lattice spacing. In the presence of activating (MAP7) and inhibitory (tau) MAPs, lattice spacing determines MAP occupancy and thereby controls kinesin-1 motility. These findings support a two-layer transport code in which MT lattice spacing directs MAP binding, and MAPs determine which motors can move along individual MT tracks.

## Introduction

Microtubules (MTs) are formed by the polymerization of α/β tubulin heterodimers into protofilaments and the association of 11-15 protofilaments laterally into a hollow cylindrical tube. These filaments serve tracks for intracellular cargos, facilitate chromosome segregation during cell division, and establish the spatial organization of organelles. The assembly and dynamics of the MT network are regulated by motors, nucleators, depolymerizers, tip-tracking proteins, and structural MAPs^1^. In turn, MTs are not merely passive tracks but active regulatory platforms that orchestrate the spatial and temporal recruitment and activity of these proteins. The molecular rules that govern how intracellular cargos are sorted to their proper destinations remain poorly understood^2^.

The tubulin-code model proposes that specific tubulin genes (also known as isotypes) and post-translational modifications (PTMs) create molecular signals that regulate MT binding and activity of motor and non-motor MAPs^3, 4^. In the human genome, there are 9 α- and 9 β-tubulin isotypes, which are differentially expressed based on cell lineage and developmental time point^5, 6^. Most of the sequence divergence between isotypes lies in the flexible, negatively charged C-terminal tails (CTTs) that project from the surface of MTs. CTTs also contain sites for many PTMs, such as enzymatic removal of the α-tubulin terminal tyrosine (detyrosination), subsequent removal of glutamate from detyrosinated α-tubulin (Δ2), and generation of branching chains of (poly)glutamate and (poly)glycine from glutamate sidechains (**Fig. 1a**)^7, 8^. The tubulin code has been shown to affect protofilament number^9^, structural features^10^, polymerization dynamics^11^, and bending rigidity of MT filaments^12, 13^. Tubulin isotypes and PTMs also regulate the recruitment and activity of structural MAPs and motors. For example, tyrosination of α-tubulin promotes recruitment of CAP-Gly-domain-containing proteins, such as CLIP170, dynactin and KIF13B, to plus-end tip complexes of MTs^14, 15^. Similarly, CTTs are directly recognized as a substrate by MT severing enzymes, and polyglutamylation promotes the MT severing activity of spastin^16^.

**Fig. 1.**
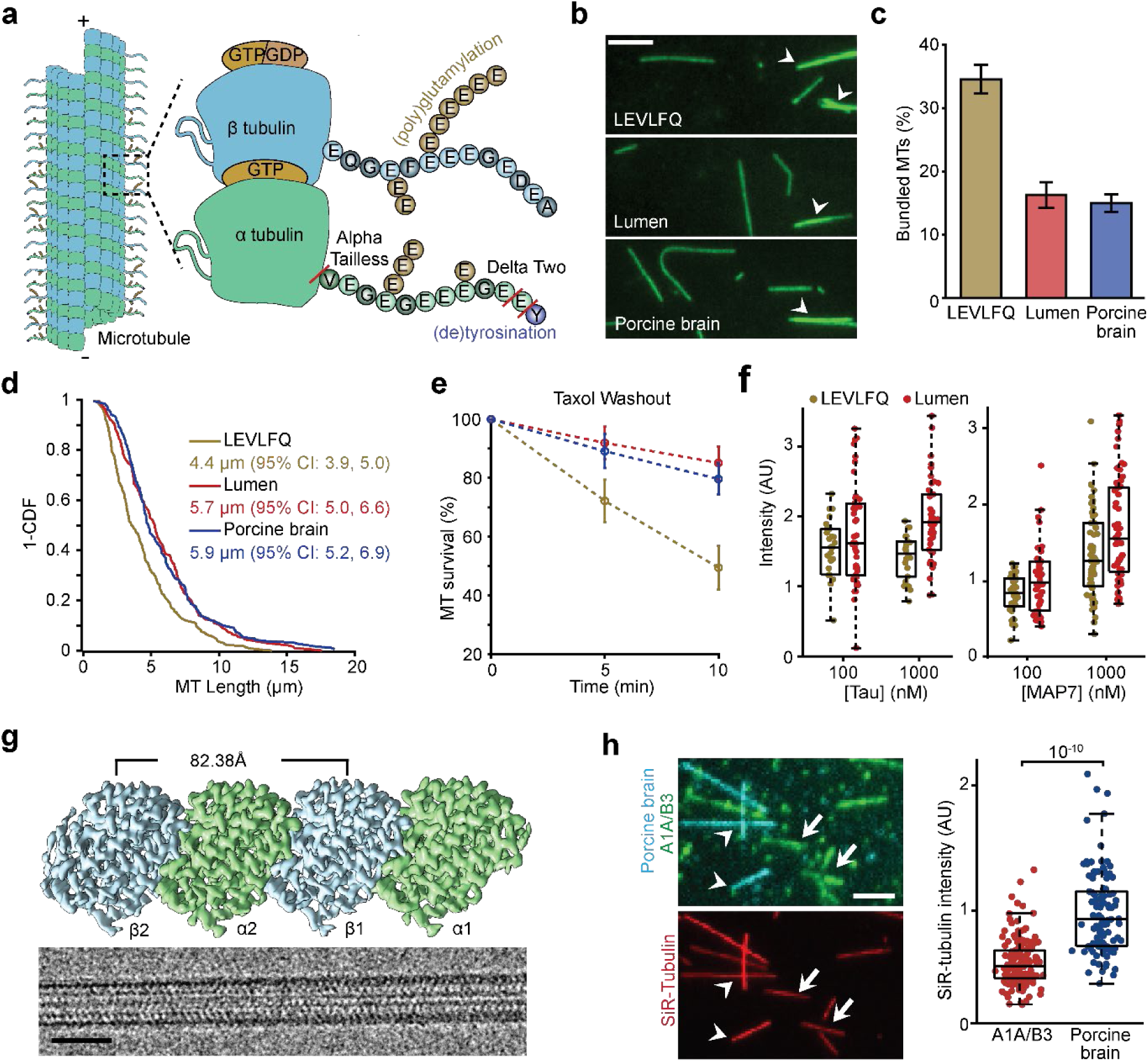
Luminal tagging of recombinant tubulin improves MT stability. a, Schematic representation of tubulin heterodimers and post-translational modifications. **b,** Representative images of MTs polymerized from α1A/β3 tubulin with a LEVLFQ protease stump on the C-terminus of β tubulin (top), from α1A/β3 tubulin with the DYKDDDK purification tag inserted inside the luminal loop of β tubulin (middle), and from porcine brain tubulin (bottom). Aberrant MTs are marked by white arrowheads. Scale bar: 5 μm. **c,** Quantification of aberrant MTs from polymerizations of different tubulins. **d,** Inverse cumulative distribution functions (1-CDF) of the lengths of Taxol-stabilized MTs polymerized from different sources. Maximum likelihood fits of each distribution to a single exponential decay constant for LEVLFQ, lumen-tagged, or porcine brain tubulin with 95% confidence intervals. **e,** Survival rate of different MTs upon the washout of the Taxol, mean survival percentage ± standard deviation for 3 technical replicates. **f,** Binding of tau and MAP7 proteins to MTs polymerized from α1A/β3 tubulin tagged at different sites. Box plots depict the median (center line), interquartile range (box), and full data range excluding outliers (whiskers). From left to right, N = 47, 42, 42, and 22 MTs for tau, and 40, 50, 56, and 62 MTs for MAP7, from two technical replicates. **g,** (Left) representative EM micrographs of lumen- tagged (top) and LEVLFQ α1A/β3 MTs (bottom). Scale bar 25nm. (Right) 3D reconstruction of lumen-tagged α1A/β3 MTs. **h,** (Left) Representative images of α1A/β3 MTs (white arrows, green) and porcine brain MTs (arrowheads, cyan) incubated with SiR-tubulin (red) in the same flow chamber. Scale bar: 5 μm. (Right) Quantification of SiR-tubulin intensity on different MTs. N = 160, 97 MTs across two technical replicates. The p-value was calculated from a mixed linear effects (MLE) model.

PTMs of tubulin CTTs also regulate MT binding and activity of other structural MAPs (such as tau, MAP2, MAP7, MAP9, and DCX) and motors in cells, although direct interactions of CTTs with these proteins could not be directly observed by cryoelectron microscopy (Cryo-EM). For example, tau preferentially binds to polyglutamylated MTs^17^. Loss of tubulin glycylation leads to the dysregulation of axonemal dyneins^18^. Kinesin-1 prefers to walk along acetylated and detyrosinated MTs^19, 20^, whereas kinesin-3 preferentially walks upon tyrosinated MT bundles^21^. MAPs and motors show substantially reduced MT binding when CTTs are cleaved enzymatically in vitro, demonstrating their functional importance^22–24^. However, in vitro reconstitution of kinesin motility on MTs with distinct PTMs did not recapitulate many of the effects seen in vivo^25^. Therefore, it remains unclear how CTTs interact with and tune the MT binding and activity of these proteins^26, 27^.

Recently, MT lattice spacing emerged as an important new dimension of the tubulin-code^28^. As tubulin incorporates into a MT, GTP at the E-site of β-tubulin is hydrolyzed to GDP, associated with compaction in the spacing between adjacent tubulin heterodimers from 83.2 Å to 81.5 Å^29, 30^. Nucleotide-dependent conformational changes are known to be critical for MT end-binding proteins to localize to the plus-end tip of MTs^31, 32^. Tau and DCX prefer compacted MT lattice whereas MAP7 prefers the expanded lattice conformation^33–35^. Kinesin-1 also prefers the extended GTP lattice^36, 37^, suggesting that lattice spacing may play a role in regulation of MAPs and motors.

The tubulin-code may also play a major role in subcellular localization patterns of MAPs, which appear to correlate with their regulatory role in intracellular transport^38^. For example, overexpression of tau inhibits kinesin-1-driven transport in axons, MAP2 promotes kinesin-3- driven cargos at the axon-initial segment, and MAP4 regulates kinesin-2 and dynein-driven transport across melanophores^39–41^. In vitro reconstitution studies demonstrated the effect many MAPs have on motors, such as tau’s inhibition and MAP7’s activation of kinesin-1^38, 42–45^, supporting that MAPs have a direct regulatory role on which motors can bind and walk along MTs. Potential crosstalk between the tubulin-code and MAP-code of MTs and how they collectively regulate motor proteins has yet to be explored.

Studying the tubulin code of MTs in vitro has been a challenge, as most studies purified tubulin from mammalian brains, which contain a diverse set of isotypes and PTMs^46^. In addition, most in vitro reconstitutions relied on Taxol, an MT stabilizing drug, which locks the lattice into an expanded conformation^47^. These obstacles have recently been overcome through mutations of tubulin-modifying enzymes and recombinant expression of isotypically pure tubulin, which provide varying levels of control over tubulin modifications^11, 17, 48, 49^, and discovery of novel drugs, such as Peloruside A (PelA), that stabilize MTs in the compacted lattice conformation^47, 50^_._

In this study, we used isotypically pure, single-PTM tubulin heterodimers to polymerize distinctly modified MTs in different lattice conformations. We tested the predictions of the tubulin- and lattice-code models on MT binding of structural MAPs with recombinantly expressed proteins in vitro. Kinesin-1 motility was reconstituted on MTs polymerized from tissue-purified or recombinant tubulin in the presence of different MAPs. These experiments directly assessed the sensitivity of MAPs and motors to tubulin CTTs and MT lattice spacing and determined whether MT tracks themselves dictate kinesin-1 motility.

## Results

### Luminal Tagging of Recombinant β3-tubulin

To study the effects of tubulin PTMs, we purified α1A/β3-tubulin heterodimers using baculovirus expression in insect cells^51–53^. This purification scheme includes an 8x histidine tag replacing the acetylation loop of α1A-tubulin and a protease sequence followed by a FLAG affinity tag on the C-terminus of β3-tubulin **(Extended Data Fig. 1a)**. Tubulin purified in this manner is isotypically pure and lacks detectable modification to the CTTs (hereafter, naïve tubulin; **Extended Data Fig. 1d-f**). We generated detyrosinated, Δ2, and α-tailless (missing residues after S439) tubulin via insertion of stop codons **(Fig. 1a).** We also polyglutamylated naïve tubulin in vitro using purified TTLL7 (see Methods)^54^.

MTs polymerized from recombinant tubulin exhibited pronounced defects compared to porcine brain tubulin, including reduced length, increased bundling, and decreased stability **(Fig. 1b-d)**. These MTs also depolymerized rapidly upon removal of Taxol **(Fig. 1e)**. Defects with recombinant MTs may be related to tagging of the β-CTT, which leaves a LEVLFQ stump after proteolytic cleavage. CTTs are vital to MT stability as proteolytic cleavage of CTTs shows pronounced bundling^55^, removal of the negatively charged residues disrupts MT dynamics^53^, and mutations to the C-terminus of β-tubulins can be lethal^56^. To mitigate the potential effects of affinity tagging, we moved the tag to the H1-S2 loop of β3 tubulin, which is solvent-exposed for soluble tubulin dimers but hidden in the MT lumen upon polymerization. Replacement of the H1– S2 loop disrupted tubulin folding, so we instead inserted a FLAG tag flanked by flexible linkers within this loop **(Extended Data Table 1, Extended Data Fig. 1e)**. Using a native β3-tubulin tail with a luminal FLAG tag, α1A/β3 MTs were no longer significantly shorter than porcine brain MTs, had a reduced proportion of bundled MTs, and depolymerized at the same rate as control **(Fig. 1b-e)**. We note that repositioning the affinity tag away from the β-CTT did not strongly affect the MT affinity of the human 2N4R isoform of tau (hereafter tau) or MAP7 **(Fig. 1f)**, suggesting that MT binding of these MAPs is not strongly sensitive to the β-CTT.

Recent reports show that α1A/β3 tubulin has a reduced affinity for Taxol and suggest that it polymerizes into MTs with a more compacted lattice^57, 58^. To test this, we imaged a Taxol-derived fluorophore, SiR-tubulin, binding to MTs polymerized from porcine brain and α1A/β3 tubulin in the same chamber **(Fig. 1h)**. We found a twofold reduction in SiR-tubulin binding to α1A/β3 MTs compared to porcine brain MTs, consistent with reduced affinity for Taxol^57, 58^. We next solved the cryo-EM structure of 14-protofilament Taxol-stabilized α1A/β3 MTs at 3.6 Å resolution (**Extended Data Fig. 2**). We find that the lattice spacing of α1A/β3 MTs (82.38 Å) is nearly identical to that of porcine brain MTs **(Fig. 1g)**. These results show that β3 tubulin exhibits lower affinity for Taxol, and similar to other tubulin isotypes, β3 tubulin remains in the expanded conformation in the presence of Taxol.

### Structural MAPs are More Sensitive to MT Lattice Spacing than Tubulin PTMs

To test whether tubulin PTMs affect MT affinity of structural MAPs, we quantified the binding of fluorescently labeled tau, MAP7, MAP9, DCX, and MAP4 to α1A/β3 and porcine brain MTs stabilized with Taxol **(Extended Data Fig. 3)**. We found MAP7 to be mostly insensitive to the tubulin content of MTs with an apparent dissociation constant between 83 and 135 nM for all MTs tested **(Fig. 2a-d)**. Tau and DCX bind to each of the differently modified α1A/β3 MTs at single-nanomolar affinity, suggesting that their MT affinity is not highly sensitive to tubulin PTMs. However, these MAPs show a nearly two orders of magnitude lower affinity for porcine brain MTs (73 ± 11 nM) when compared to α1A/β3 MTs (1.7 ± 0.5 nM; **Fig. 2d, Extended Data Fig. 3)**.

**Fig. 2.**
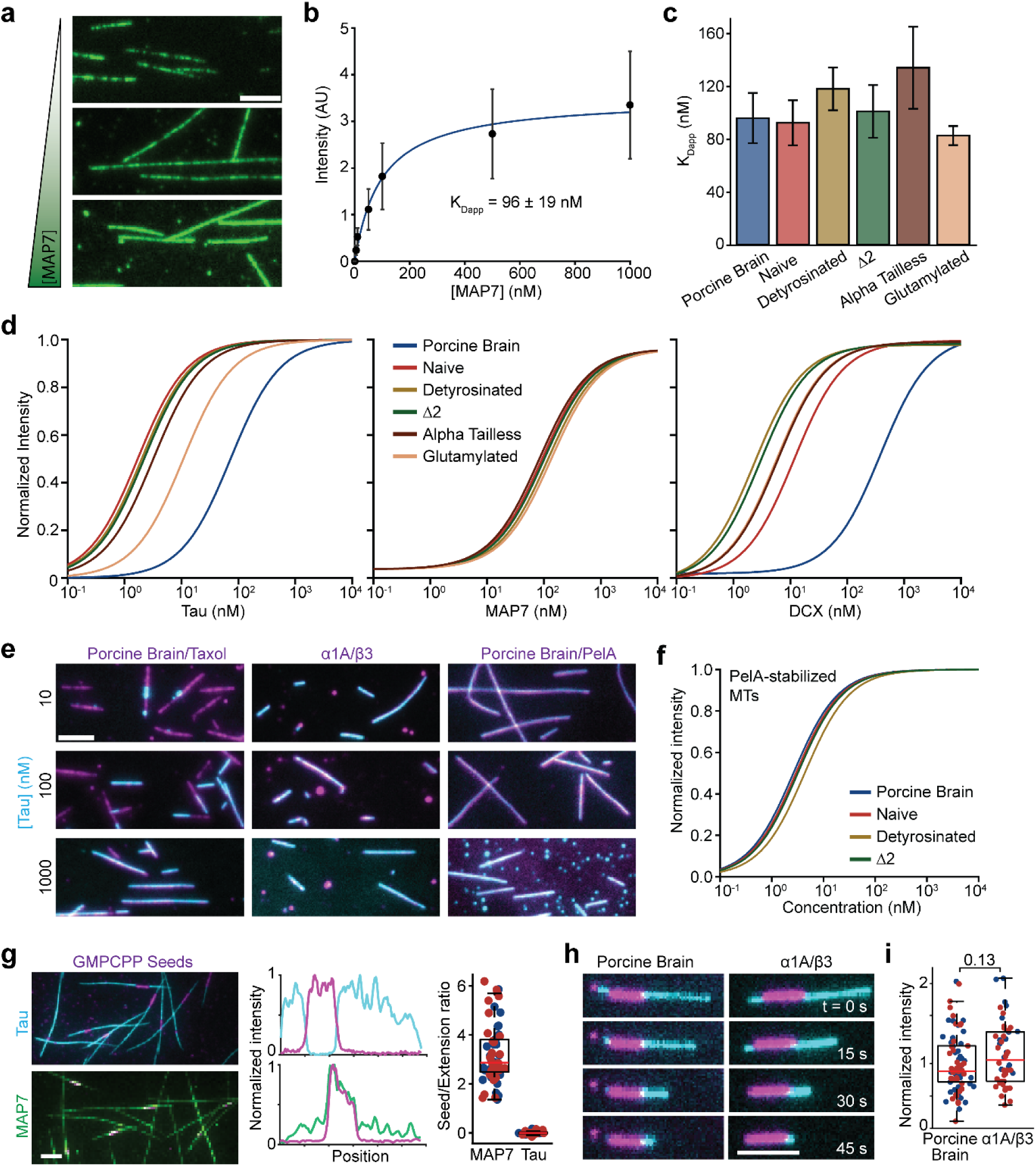
MAP binding is differentially regulated by MT lattice spacing. a, Representative images of MAP7 binding to porcine brain MTs at increasing concentrations (10, 100, 1000 nM), scale bar 5 μm. **b,** The MAP7 intensity on MTs as a function of MAP7 concentration. A fit to a Hill equation with n = 1 (solid blue curve) reveals apparent K_D_ = 96 ± 19 nM (mean ± S.E. of fit). **c,** Apparent K_D_ values of MAP7 binding to different MTs (mean ± S.E.). **d,** Hill equation curve fits for MAP7, tau, and DCX binding to differently modified Taxol-stabilized MTs. **e,** Representative images of tau (cyan) binding to MTs (magenta), scale bar: 5 μm. **f,** Curve fits for tau binding to differently modified PelA-stabilized MTs. **g,** (Left) Representative images of 100 nM tau (cyan) and MAP7 (green) binding to dynamically growing MTs and GMPCPP seeds (magenta). (Right) Representative line scan intensities of tau (cyan) and MAP7 (green) compared to GMPCPP seed intensities (magenta). (Bottom) Ratios of fluorescence intensities of MAPs localized to MT extensions over MAPs localized to GMPCPP seeds. Mean ratio of seed to extension intensity is fold change is 3.2 ± 1.2 (mean ± S.D.) for MAP7 and 0.002 ± 0.05 for tau. N = 60 and 56 MTs across two technical replicates. Colors denote replicates. **h,** Representative images of tau (cyan) binding to GDP-MT extensions from GMPCPP seeds (magenta) after washout of free tubulin. **i,** Quantification of tau intensity on GDP-MTs 15 s after washing out free tubulin. N = 78 and 50 MTs across two technical replicates. Colors denote replicates. The p- value was calculated from an MLE model.

We next explored whether the differences in MT binding affinity of tau for α1A/β3 and porcine brain MTs were due to the stabilization of the expanded MT conformation by Taxol. Because tau has higher affinity for the compacted MT lattice^35, 59, 60^ , we first quantified tau binding to GDP MTs. We grew compacted GDP-lattice extensions from extended GMPCPP-stabilized MT seeds in the presence of soluble tubulin **(Fig. 2g)**. Strikingly, tau binds robustly to the GDP lattice extensions with only a 0.1% relative intensity on GMPCPP seeds **(Fig. 2g)**. Interestingly, we observed that tau intensity tapers off towards the plus-end of dynamic MTs, consistent with a higher proportion of expanded, GTP-tubulin **(Fig. 2g)**. We found the opposite behavior for MAP7, which shows a 3.2-fold enrichment on the GMPCPP seeds when compared to the GDP lattice extensions. This is consistent with emerging research suggesting that MAP7 prefers to bind to expanded MT lattices in vivo (**Extended Data Fig. 4**)^33, 61^. To eliminate potential effects of soluble tubulin, we grew the GDP lattice extensions from the seeds, removed free tubulin while introducing tau to the chamber, and quantified tau intensity on the MTs while they quickly depolymerized **(Fig. 2h, i)**. We found that tau binds similarly to GDP-lattice MTs polymerized from porcine brain or α1A/β3 tubulin.

Finally, we tested tau-MT binding using PelA^50^, which stabilizes MTs in a compacted lattice spacing^47^. Tau had similar (2.5 - 4.5 nM) affinity to bind all PelA-stabilized MTs polymerized from brain vs. recombinant tubulin **(Fig. 2f, Extended Data Fig. 5)**. This affinity of tau was comparable to Taxol-stabilized α1A/β3 MTs **(Fig. 2b, Extended Data Fig. 3)**, raising the possibility that tau can outcompete Taxol from recombinant MTs because of low affinity of β3 tubulin for Taxol, whereas higher tau concentrations are required to compact the lattice of pig brain MTs. Collectively, these results show that MAP binding to MTs is not strongly sensitive to the tubulin PTMs we tested and that MT lattice spacing is a larger driver of tau and MAP7 binding to MTs.

### Tau Reads MT Lattice Spacing Through the MT Binding Domain

We next turned our attention to understanding how tau binding changes between compacted and expanded MT lattices. Previous studies have shown that, at low concentrations, tau weakly interacts with and diffuses along Taxol-stabilized MTs. At higher concentrations, tau molecules interact with each other through liquid-liquid phase separation (LLPS) and form envelopes on these MTs^42, 62^. Inside the envelopes, tau stably binds to the MT and exhibits little diffusion. These observations suggested that single tau molecules have low affinity for MTs, and interactions between many tau molecules increase the avidity for more stable binding to the MT. Tau envelope formation also reduced SiR tubulin binding to those regions^35^, suggesting that envelope formation of tau compacts the MT lattice, raising the alternative possibility that MT compaction stabilizes tau binding to the MT.

Previous reports had shown that the tau construct that contains the MT binding domain but lacks the projection domains (tau-MTBD) and hyper-phosphorylated full-length tau have a greatly reduced affinity for MTs compared to wild-type, full-length tau (FL-tau)^62, 63^. Lower apparent MT affinity of these constructs has been attributed to their inability to efficiently interact with each other through LLPS. We attempted binding assays of tau-MTBD and tau-14E (a phosphomimic mutant with 14 non-MTBD sites mutated to glutamate) to Taxol-stabilized MTs and confirmed that these tau variants show very little binding **(Fig. 3a)**^64^. We only found these variants binding as puncta on curved regions of MTs, consistent with tau’s preference to bind MT bends^62^ In contrast to Taxol-stabilized MTs, we observed that tau-MTBD and tau-14E bind uniformly along the entire length of PelA-stabilized MTs **(Fig. 3a)**. The apparent dissociation constants of tau-MTBD (238 ± 56 nM) and tau-14E (185 ± 43 nM) are similar to FL-tau binding to Taxol MTs, but two orders of magnitude less than FL-tau binding to PelA MTs **(Fig. 3b, Extended Data Fig. 5)**. Fits of their binding curves revealed a larger Hill coefficient for FL-tau (1.4 ± 0.4, mean ± S.E. of the fit) than for tau-MTBD (1.2 ± 0.1) and tau-14E (1.1 ± 0.1), agreeing with the hypothesis on the loss of cooperativity from favorable tau-tau interactions due to phosphorylation^63^.

**Fig. 3.**
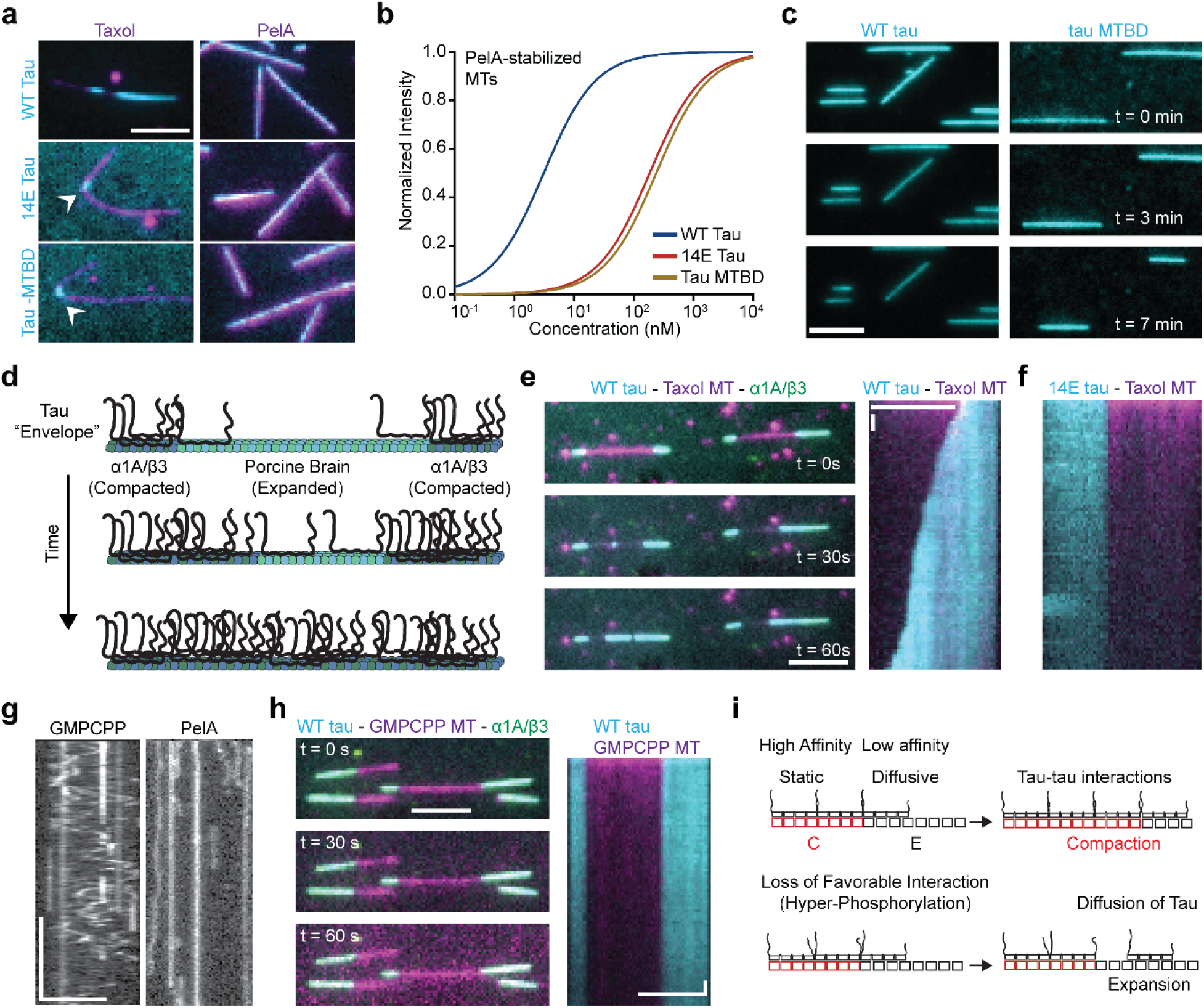
**Tau separately reads and writes the MT lattice state**. **a,** Representative images of 100 nM tau variants (cyan) binding to Taxol and PelA stabilized MTs (magenta), scale bar 5 μm. White arrowheads point to tau enrichment on curved MTs. **b,** Hill equation curve fits for Tau variants binding to PelA MTs. **c,** Representative images of tau localized to MTs after washing tau out of solution, scale bar 5 μm. **d,** Schematic for the condensate growth assay along MTs. **e,** (Left) Representative images of tau condensate growth over time from α1A/β3 MT ends, scale bar 5 μm. (Right) Representative kymograph of tau condensate growth from α1A/β3 MT caps into Taxol-stabilized lattice, scale bars: 5 μm and 10 s. **f,** Representative kymograph of 14E tau growth from compacted MT caps into Taxol-stabilized lattice**. g,** Single-molecule diffusion of different tau variants on GMPCPP- and PelA-stabilized lattices. **h,** Representative images of tau condensate growth over time from compacted MT ends (Left), scale bars: 5 μm and 10 s. (Right) Representative kymograph of abolished tau condensate growth from compacted MT caps into GMPCPP-stabilized lattice. **i,** Schematic representation of tau compacting the MT lattice.

After determining that tau-MTBD recognizes and binds to the compacted lattice, we attempted to determine the off-rate of tau variants to PelA MTs. We anticipated that wild-type tau would stay tightly bound to MTs through multiple tau-tau and tau-MT interactions, but tau-MTBD would quickly dissociate when removed from solution. However, we found that both wild-type tau and tau-MTBD remain stably bound to PelA MTs over the course of 7 min after washout **(Fig. 3c)**. These findings suggest that tau condensates resist dissociation from the MT through tight tau-MT interactions rather than tau-tau interactions.

Previously, it was found that tau diffuses quickly on MTs outside of tau condensates and remains stationary inside condensates^62^. This was concluded to be due to tau-tau interactions holding tau in place, as in a liquid phase. We suspected that the loss of tau diffusion in condensates may not be due to tau-tau interactions, but rather to changing tau-MT interactions at different lattice compactions. We tested tau diffusion on differently stabilized MTs at single-molecule concentrations, to watch tau motility without confounding tau-tau interactions. We find that single tau molecules diffuse very quickly on GMPCPP-stabilized lattices, but are stationary on PelA lattices, even in the absence of other tau molecules **(Fig. 3g)**. This result shows that tau stably binds to the MT lattice inside the envelopes by strongly interacting with the compacted lattice, not through a network of tau-tau interactions.

### Tau Writes MT Lattice Spacing Through the MTBD Flanking Regions

We hypothesized that the reason tau-MTBD and tau-14E show little to no MT binding in previous assays is that these constructs are deficient in compaction of the Taxol-stabilized MT lattice. To test this, we generated short α1A/β3 MT extensions on Taxol-stabilized porcine brain MT seeds (see Methods; **Fig. 3d)**. We then flowed tau into the chamber, which quickly binds to the α1A/β3 extensions but not to the seeds. FL-tau envelopes gradually grew from the α1A/β3 extensions into the porcine brain MT seeds **(Fig. 3e)**. Unlike FL-tau, we find that tau-14E binds quickly to the compacted MT extensions but does not grow condensates into the Taxol-stabilized lattice: confirming its reduced ability to compact the lattice^65^ **(Fig. 3f)**. When we grow α1A/β3 extensions from GMPCPP-stabilized seeds in the presence of Taxol, we find that both tau-14E and FL-tau cannot grow condensates into seeds, as the GMPCPP lattice does not hydrolyze to GDP-tubulin **(Fig. 3h)**. Without hydrolysis to GDP-tubulin, GMPCPP seeds cannot transition to the compacted lattice state, locking the MT into low affinity for tau.

These findings suggest that tau-tau interactions on the expanded MT lattice facilitate tau-induced compaction of the lattice. In this model, single tau molecules transiently interact with and diffuse along an expanded MT lattice **(Fig. 3i)**. Tau-tau interactions on the expanded lattice reduce the dissociation rate of individual tau molecules from the MT and increase their local concentration on the MT. Tau binding under these conditions locally compacts the MT lattice, which stabilizes MT binding of individual tau molecules inside the envelope. Diffusive tau molecules on nearby uncompacted lattice sites interact with tau molecules at the boundary of these envelopes, which induces further compaction of the MT lattice and the growth of tau condensates. Tau-induced MT compaction leads to their cooperative binding to MTs. Interestingly, phosphorylation of tau’s projection domains negatively influences the ability of tau molecules to interact with each other and hence acts as a potentiometer that reduces tau’s capacity to compact the MT lattice.

### MAP Competition for the MT is Sensitive to Lattice Compaction and Detects MT Curvature

Since we observed large changes in MT affinity for MAPs, we next questioned if these differences functionally alter the decoration of MTs in a complex mixture. To test this, we performed two-MAP competition assays with tau and MAP7 **(Fig. 4a)**. We find that MAP7 excludes tau on Taxol-stabilized porcine brain MTs, except at curved regions, consistent with previous reports **(Fig. 4b)**^66^. Remarkably, on all differently modified α1A/β3 MTs, tau consistently outcompeted MAP7 for MT occupancy, except for α tailless MTs, where we observed co-binding of MAP7 and tau **(Fig. 4b)**. This preference for tau binding is consistent with the compaction of α1A/β3 MTs, and the co-decoration of tau and MAP7 on α tailless MTs may be related to potential docking of CTTs upon lattice compaction^67^. When we performed MAP7 vs tau competitions on porcine brain MTs stabilized with PelA, we found that MAP7 no longer excludes tau **(Fig. 4b)**. Interestingly, we found MAP7 to be excluded on PelA MTs, except at curved regions **(Fig. 4a)**. We performed additional competition assays with other two-MAP pairs and found that decoration switches when one MAP is highly sensitive to lattice spacing (such as tau; **Extended Data Fig. 6)**. We did not find MAP decoration switching between the different physiologically relevant PTMs tested.

**Fig. 4.**
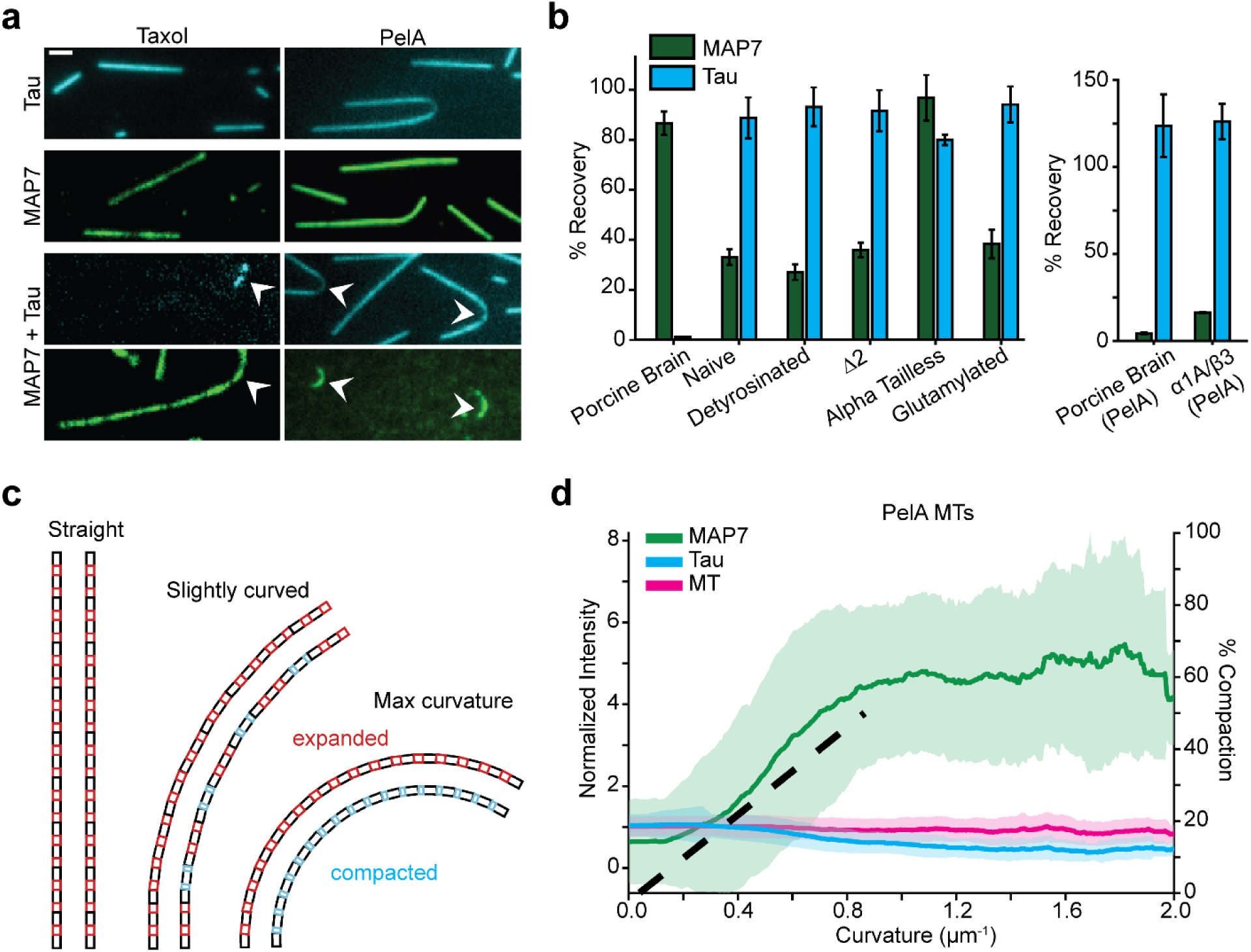
MT lattice spacing directs MAP competition and marks MT curvature. a, Representative images of a two-MAP competition assays between MAP7 and tau. White arrowheads point to tau and MAP7 enrichment at curved MTs. **b,** Quantified competitions between tau and MAP7 on Taxol-MTs (left) and PelA-MTs (right). Bars represent mean fluorescence recovery of three replicates (± SD). **c,** Representative schematic of lattice compaction due to MT curvature. As Taxol-MTs bend, inner protofilaments become compacted and outer protofilaments stay expanded. **d,** Fluorescence intensity as a function of MT curvature for MAP7 and tau on PelA MTs. MAP7 in green, tau in cyan, MT in magenta. Thick bars represent mean fold-increase over average MT intensity, and shaded regions denote standard error. Black dashed line is the function of compacted lattice sites as a MT becomes more bent. Data from 331 curved MTs across three technical replicates.

In our competition assays, we found specific enrichment of the less competitive MAP at the curved regions of MTs **(Fig. 4a)**. At bent regions, MTs have heterogeneous lattice spacing regardless of stabilization method; the interior of the bend would have compacted protofilaments, whereas the exterior of the bend would be expanded **(Fig. 4c)**. Assuming that tubulin behaves as a two-state system with lattice spacings for GDP- and GMPCPP-MTs, we approximate that a MT would need a radius of curvature of 1.2 μm to have a fully compacted inner-protofilament and fully expanded outer-protofilament^30^. We analyzed the fold-enrichment of tau and MAP7 as a function of MT curvature from our assays (**Fig. 4d**, see Methods). We find that on PelA-MTs, MAP7 is enriched at bent regions and reaches a plateau at bends with a radius of curvature of 1.3 μm **(Fig. 4d, left)**. The increase in MAP7 decoration at curved regions is concomitant with a decrease in tau decoration. The smooth increase of MAP7 enrichment and tau depletion aligns well with the function of how many lattice sites are compacted as the MT becomes more bent (dashed black line, **Fig. 4d**). We find the opposite relationship on Taxol-MTs, where tau is enriched in curved regions as previously described **(Extended Data Fig. 7)**^62^. These results indicate that MT curvature induces lattice compaction and extension, which may be a mechanism for MAPs and motors to sense bent or buckling MTs, such as in the beating flagellum or contracting myocardiocytes.

### Kinesin-1 Motility is Insensitive to Tubulin PTMs, but is Tuned by MT Lattice Spacing

To assay kinesin-1’s activity on MTs polymerized from different tubulins, we grew long Taxol stabilized MT extensions from porcine brain GMPCPP seeds in our imaging chambers **(Fig. 5a)**. We found that kinesin-1 has a 50% reduced run frequency on α1A/β3 MTs compared to porcine brain MTs **(Fig. 5b)**. There was little to no change in kinesin-1’s run frequency on differently modified α1A/β3 MTs, suggesting that kinesin-1 is more sensitive to the lattice spacing of the MT than the tubulin PTMs tested **(Fig. 5b)**. Notably, the lattice preference of kinesin-1 was similar to its activating MAP, MAP7, which also preferred binding to GMPCPP seeds and opposite to its inhibitory MAP, tau, which strongly preferred the compacted lattice.

**Fig. 5.**
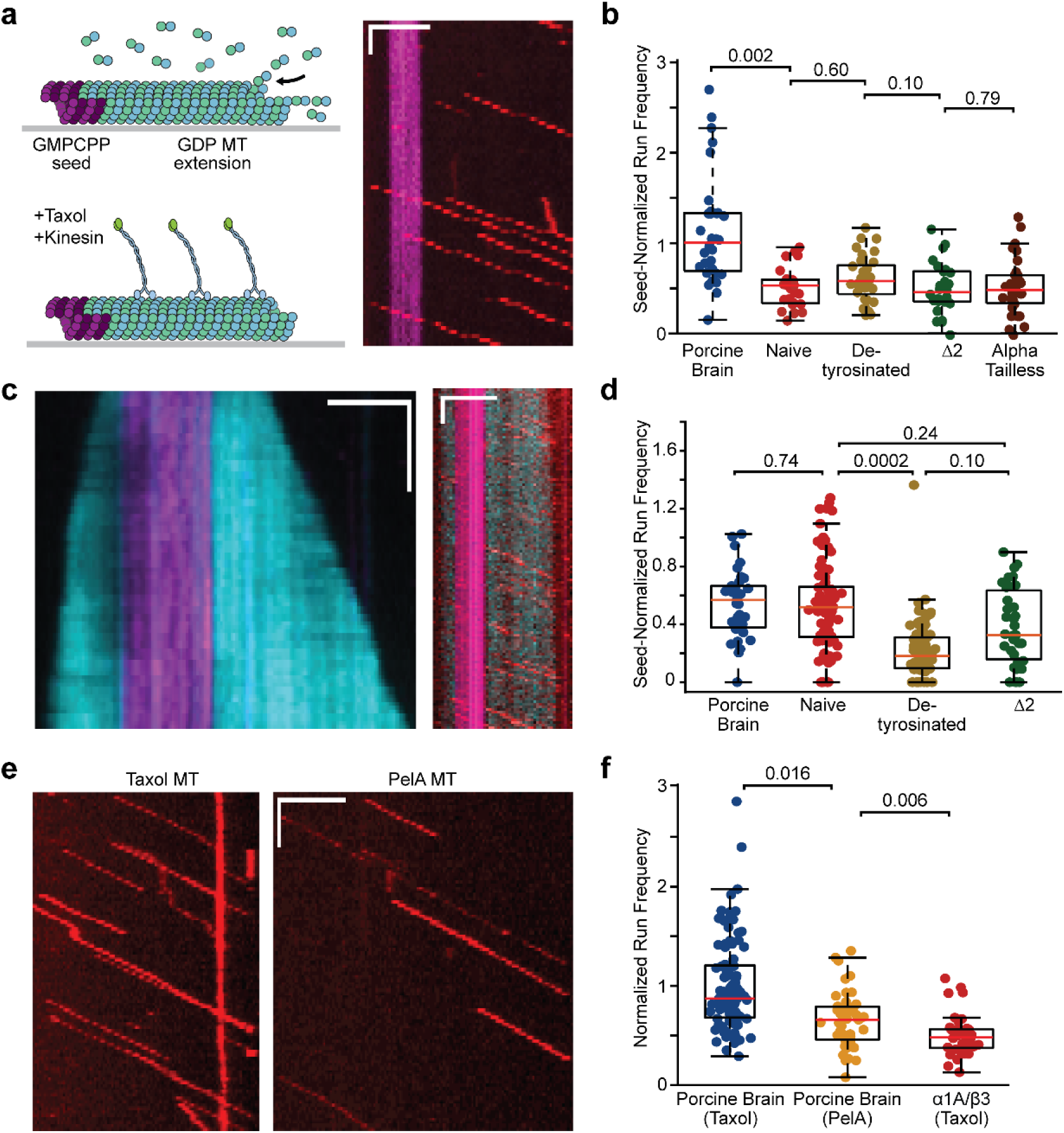
MT lattice spacing, but not PTMs, tunes kinesin-1 motility. a, (Left) Schematic representation of preparation of long MTs for in vitro motility assays. (Right)Representative kymograph of kinesin-1 motility on porcine brain MTs, scale bars 5 μm and 10 s. **b,** Quantified run frequencies on differently made MTs normalized to kinesin run frequency on GMPCPP seeds in the same chamber. N = 32, 36, 37, 32, 31 MTs across three technical replicates. The p-values were calculated from an MLE model. **c,** (Top) Representative schematic of dynamic MT kinesin motility assay. (Bottom) Representative kymograph of MT extension, scale bars 5 μm and 1 min. (Right) Representative kymograph of kinesin-1 activity on dynamically growing MTs, scale bars 5 μm and 10 s. **d,** Kinesin-1 run frequencies on dynamically growing MTs grown from different tubulins normalized to the run frequency on GMPCPP seeds in each experiment. N = 35, 81, 100, 35 MTs across three technical replicates. P-values from an MLE model. **e,** Representative kymographs of kinesin-1 motility on Taxol (left) and PelA (right) stabilized MTs. **f,** Run frequency quantification of kinesin-1 on different MTs, normalized to MT length and imaging time. N = 96, 43, 37 MTs across three technical replicates. P-values calculated from an MLE model.

We performed kinesin-1 motility assays in dynamic MT conditions where MTs were actively polymerizing **(Fig. 5c)**. We noticed that dynamically growing MTs had large variations in kinesin-1 motility, suggesting that the tubulin in solution has a large impact on kinesin motility **(Extended Data Fig. 8a)**. We also found large differences in kinesin-1 motility on porcine brain GMPCPP seeds depending on which tubulin is in solution **(Extended Data Fig. 8b)**. To account for the sequestration of kinesin-1 by soluble tubulin, we normalized the run frequency of each MT to the average run frequency of kinesin-1 on the GMPCPP seeds in each experiment as an internal standard. We find that MTs polymerized from porcine brain or unmodified α1A/β3 tubulin do not significantly differ from each other, and that all GDP MT extensions have a reduced kinesin-1 run frequency when compared to GMPCPP seeds **(Fig. 5d)**. We also note a modest reduction in kinesin-1 run frequency on detyrosinated α1A/β3 MTs under these conditions **(Fig. 5d)**.

To eliminate the effect of free tubulin from the solution while studying motility on compacted MTs, we used PelA-stabilized porcine brain MTs as a model for compacted MT lattice. We find that kinesin-1 has a 34% reduced run frequency on PelA-stabilized MTs as compared to Taxol-stabilized MTs, consistent with its preference for expanded MT lattices **(Fig. 5f)**^57^. This reduced run frequency on PelA-stabilized porcine brain MTs is closer to kinesin-1’s activity on α1A/β3 MTs **(Fig 5f)**. Collectively, our results revealed that kinesin-1 is more sensitive to MT lattice spacing than tubulin PTMs, but the effects are modest.

### MT Lattice Spacing Directs Kinesin-1 Motility Through Competitive MAP Decoration

After determining that MTs can direct MAP decoration, we wanted to test how changing MAP decoration could impact motor activity. Previous in vitro studies concluded that tau condensates are a potent inhibitor of kinesin-1 motility, whereas MAP7 increases the number of kinesins that land and walk on the MT^42, 44, 45^. We suspected that the lattice state can play a major role in the MAP-mediated regulation of kinesin-1. We performed kinesin-1 motility assays in the presence of MAP7 and tau on Taxol-stabilized MTs made of either porcine brain or α1A/β3 tubulin **(Fig. 6a)**. On porcine brain MTs, MAP7 excluded tau and enhanced kinesin-1 run frequency three-fold. On α1A/β3 MTs, tau excluded MAP7 and inhibited kinesin-1 from walking, except on GMPCPP seeds. On GMPCPP seeds, tau cannot compact the MT and form condensates, so kinesin-1 is able to land and walk until it encounters a tau condensate at the border of α1A/β3 MTs **(Fig. 6a, b)**. Repeating this experiment with PelA-stabilized porcine brain MTs, we again find tight tau binding and an abolishment of kinesin-1 motility **(Fig. 6c, d)**. Consistent with our findings on MAP decoration and previous work on MAP-motor interactions, these results show that the lattice spacing of MTs controls kinesin-1 activity through MAP recruitment in a complex environment.

**Fig. 6.**
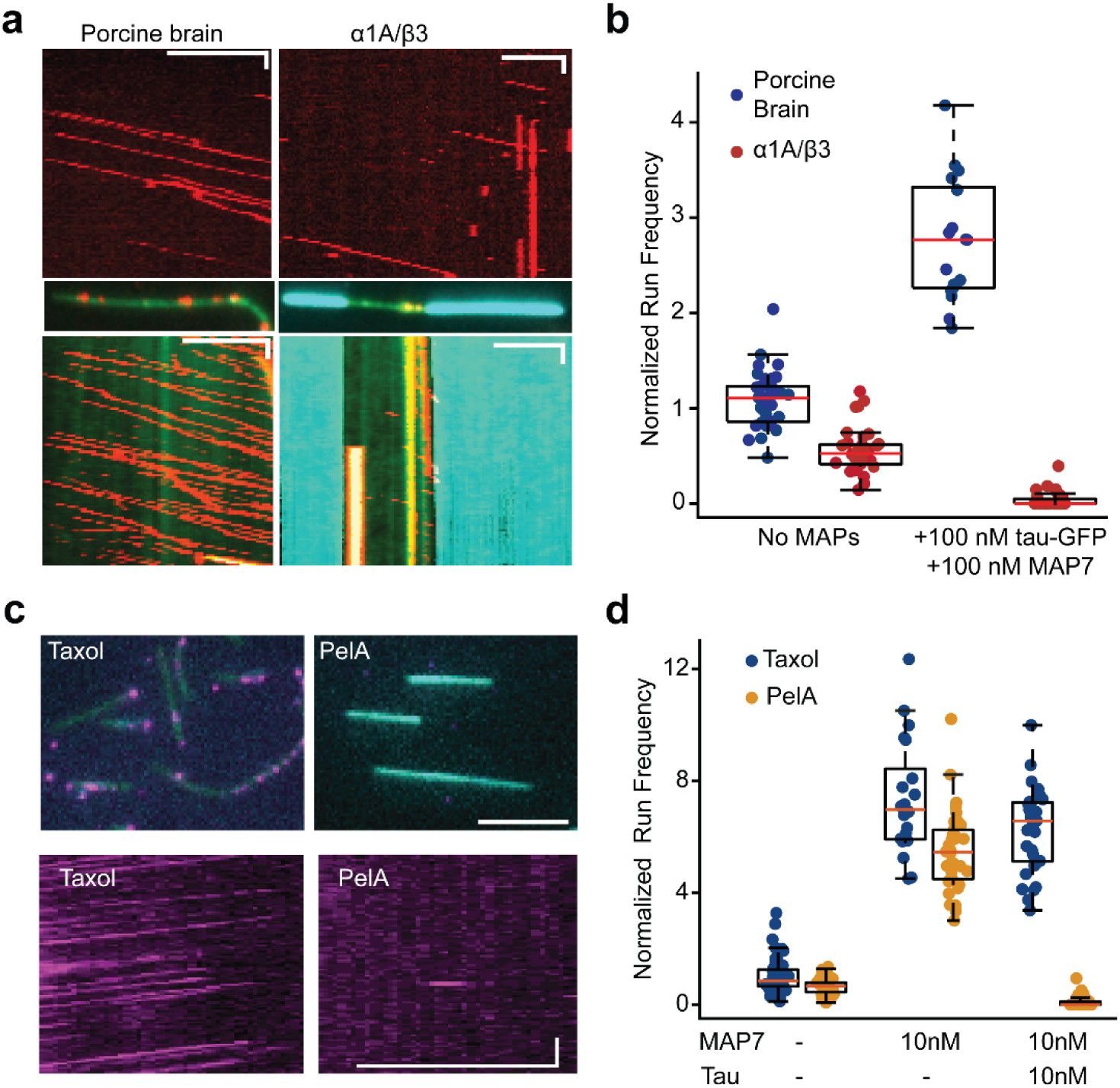
The lattice code directs MT Binding of MAPs that regulate kinesin-1 motility. a, Top, representative kymographs of kinesin-1motility on MTs without MAPs. Middle, representative images of MTs in the presence of 2 nM kinesin-1 (red), 100 nM MAP7 (green), and 100 nM tau (cyan). Bottom, representative kymographs of kinesin-1 motility on MTs in a MAP7 and tau mixture. Scale bars 5 μm and 10 s. **b,** Run frequency of kinesin-1 on naked and MAP-decorated MTs. N = 34, 37, 17, 35 MTs across two technical replicates. **c,** Left, representative images of kinesin-1 (magenta) and tau (cyan) on MTs (green). Right, representative kymographs of kinesin- 1 motility on differently stabilized MTs in 10 nM MAP7 and 10 nM tau solution. Scale bars 5 μm and 10 s. **d,** Kinesin-1 run frequency on Taxol- or PelA-stabilized MTs in the presence of different MAPs. N = 83, 43, 21, 34, 33, 53 MTs across two technical replicates.

## Discussion

Using recombinant α1A/β3 MTs, we tested MAP and motor activity against different PTMs and found the largest changes were between α1A/β3 MTs and porcine brain MTs, rather than between PTMs. These results suggested that the proteins we tested were more sensitive to the tubulin isotype than PTMs on CTTs. Using Taxol and PelA, we could replicate the changes in MAP binding between α1A/β3 and porcine brain tubulin, consistent with the different lattice spacings for Taxol- and PelA-MTs. Collectively, these results demonstrate that MT lattice spacing is the primary driver of MAP-MT interactions, more so than tubulin PTMs for the MAPs we tested.

Recombinant tubulin serves as an ideal platform for studying the tubulin code, as it gives control over each PTM and isotype to be tested in vitro. Tagging strategies should be carefully considered in future studies, as tags that lie in the MT lumen, such as in this study, would presumably interfere with MT inner proteins (MIPs). As we show, the LEVLFQ protease stump on the C-terminus of β-tubulin causes aberrant MT bundling, but potentially different polymerization conditions or different protease sequences may mitigate these artifacts.

Differences between compacted and expanded MT lattices appear to be small on the tubulin heterodimer scale. However, many MAP proteins bind MT filaments by contacting more than a single dimer. Tau has four MT binding repeats, each long enough to span a tubulin dimer. When propagated along four dimers, these small changes can accumulate enough to potentially limit the binding of multiple repeats and cause dramatic changes in affinity. We suspect that conformational changes within the tubulin subunits upon compaction drive these changes in affinity.

We determined that the tau-MTBD reads the lattice state of the MT, binding at nanomolar concentrations to the compacted lattice, but not to the expanded lattice. Tau’s capacity to compact the lattice depends on the MTBD flanking regions and is reduced by physiologically relevant phosphomimic mutations. In contrast, Taxol stabilizes an expanded lattice state, effectively acting as an antagonist to tau’s compaction activity. In vitro, this opposition creates a competition between lattice expanders and compactors, generating heterogeneous MT populations with distinct tau condensates. We note that to compact the lattice in the presence of 10µM Taxol, tau must be present at high concentration (100 – 1000 nM), whereas it acts as a potent kinesin-1 inhibitor at much lower concentrations (10 nM) on pre-compacted MTs. Together, these findings support a model in which competition between lattice expanders and compactors produces heterogeneous MT lattices that can be selectively read by MAPs, providing a potential mechanism for generating specialized MT tracks in cells.

Our results have important functional implications for MT-based transport regulation. We found modest effects on kinesin-1 motility based on either the PTMs or the lattice perturbations we tested. However, by controlling the decoration of MAPs, MT lattice spacing strikingly directs the binding of tau or MAP7. In the presence of MAPs, kinesin-1 motility was either upregulated by the presence of MAP7 or abolished by the presence of tau, depending on the MT lattice state. Recently published MT structures inside cells show that MT lattice spacing changes along neuronal differentiation^68^, but it remains unclear which effectors are most responsible for driving these changes. We propose that MT lattice spacing may act as a primary regulator for the recruitment of MAPs, which recruit or inhibit modifying enzymes and motors to distinguish intracellular transport lanes and other MT functions^33, 69^.

## Materials and Methods

### Protein Purification

Porcine brain tubulin was prepared by two cycles of polymerization and depolymerization in 1 M PIPES buffer, as previously described^46^. Briefly, brain lysate was clarified by spinning at 8,000 rpm in a Beckman 8.1000 rotor for 1 h. MTs were polymerized by mixing equal volumes of clarified lysate, High Molarity PIPES Buffer (HMPB: 1M PIPES, 10mM MgCl_2_, 20mM EGTA pH=6.9), and glycerol, then adding 1.5 mM ATP and 0.5 mM GTP. Polymerizing MTs were kept at 37 °C with shaking in a warm water bath for 1 h. MTs were pelleted at 54,000 rcf for 1 h in a Type 19 rotor at 37 °C. Pelleted MTs were washed with warm HMPB, then resuspended in cold depolymerization buffer and kept on ice for 15 min. Depolymerized MTs were centrifuged in a Type 45Ti rotor at 235,000 rcf for 30 min at 4 °C. MTs were again polymerized by mixing equal volumes of clarified supernatant, HMPB, and glycerol, then adding 1.5 mM ATP and 0.5 mM GTP. Polymerizing MTs were kept at 37 °C with shaking in a warm water bath for 30 min. MTs were pelleted in a Type 45Ti rotor at 235,000 rcf for 30 min at 37 °C. MT pellets were washed with warm BRB80, then resuspended in a minimal volume of cold BRB80 and incubated on ice for 15 min. The resuspended MTs were pelleted in a TLA 100.4 rotor at 541,000 rcf for 10 min. The supernatant was mixed by pipetting, diluted to 20 mg/mL tubulin with BRB80, and snap frozen in LN2.

Recombinant human α1A/β3 tubulin heterodimers were expressed in SF9 cells, as previously described^11, 51, 52^. Cells were lysed by two passes in an LM-20 Microfluidizer at 12,000 psi in BRB80 supplemented with 1 mM PMSF, 0.5 mM ATP, 0.5 mM GTP, Roche protease inhibitor tablets, and 25 mM imidazole. Lysate was clarified in a Type 45Ti rotor at 235,000 rcf for 45 min at 4 °C. Clarified lysate was incubated with HisPur Ni-NTA (Thermo) resin for 45 min at 4 °C to bind tubulin to the column, then washed with 5 column volumes of lysis buffer supplemented with 500 mM KCl. Tubulin dimers were eluted using cold BRB80 supplemented with 500 mM KCl and 500 mM imidazole. Ni-NTA eluate was then diluted 2-fold with BRB80 and incubated with Anti-DYKDDDDK Affinity resin (Pierce) for 1 h at 4 C. The resin was washed with 10 column volumes of BRB80, and resuspended in BRB80 supplemented with 30 μM GTP, 1.5 mg/mL 3x FLAG peptide, and 400 U HRV 3C protease (GenScript). After incubation at 4 °C for 16 h, the elution from the anti-FLAG resin was cleaned up on a 1 mL HiTrap Q HP IEX column (Cytiva). The tubulin was buffer exchanged into BRB80 supplemented with 10% glycerol, concentrated to 4.5 mg/mL, and snap frozen in liquid nitrogen.

To move the affinity tag away from the β tubulin C-terminus, we inserted a double stop codon (ATTACT) after K450 and a FLAG tag flanked by flexible linkers (GGGSDYKDDDDKGGGGSG) in between G38 and D39 of β tubulin. For tubulin with the FLAG tag in the β tubulin luminal loop, the Ni-NTA eluate was bound to ANTI-FLAG M2 Affinity Gel (Sigma-Aldrich) and eluted with 0.15 mg/mL 3xFLAG peptide (APExBio), without the protease step.

Strep-sfGFP-2N4R Tau (tau), 6xHis-DCX-eGFP-strep (DCX), strep-sfGFP-MAP9 (MAP9) were expressed in Rosetta 2 E. Coli, grown to 0.6 OD_600_ in Terrific Broth media at 37 °C. The culture was induced with 1 mM IPTG at 16 °C, incubated for 16 h with shaking at 180 rpm, and then harvested by centrifugation at 3,985 rcf. Cells were lysed by tip sonication on ice for 4 min active time in lysis buffer (50 mM Tris-HCl pH 8.0, 50 mM KAc, 2 mM MgCl2, 10% glycerol) with 0.5 mg/mL lysozyme added. Lysate was clarified in a Type 45Ti rotor at 235,000 rcf for 45 min at 4 °C. Clarified lysate was incubated with Streptactin XT 4flow resin (IBA Life Sciences) for 1 h at 4 C with rolling. Resin was collected in a Bio-Rad disposable column and washed with 10 column volumes of lysis buffer. Protein was then eluted with SFP buffer (25 mM HEPES pH 7.4, 300 mM KCl, 10 mM MgCl2, 1 mM EGTA, 10% glycerol) supplemented with 100 mM biotin. Tau and DCX were further purified on a 1 mL HiTrap S IEX column (Cytiva), and MAP9 on a Superose 6 Increase 10/300 GL size exclusion column (Cytiva). Proteins were exchanged into SFP buffer, concentrated, and snap frozen in liquid nitrogen.

Strep-SNAP-MAP4 (MAP4) was expressed in SF9 cells via baculovirus. Cells containing MAP4 were lysed by dounce homogenizer on ice, and purified using the methods described for tau, DCX, and MAP9.

ZZ-TEV-ybbR-MAP7 (MAP7) and ZZ-TEV-K560-GFP-SNAP (Kinesin) were expressed in SF9 cells via baculovirus using the methods described for α1A/β3 tubulin. Cells were lysed by dounce homogenizer on ice, lysate clarified by a Type 45Ti rotor, and lysate bound to IgG resin at 4 °C for 1 h. Proteins were eluted from the IgG resin by incubation with TEV protease at room temperature for 1 h in SFP buffer. The eluate was then concentrated and purified with a Superose 6 Increase 10/300 GL size exclusion column. Peak fractions were pooled, concentrated, and snap frozen in LN2.

TTLL7 was prepared, as previously described^54^. GST-TEV-TTLL7(1-518) (TTLL7) was expressed in Rosetta 2 E. Coli, grown to 0.6 OD_600_ in Terrific Broth media at 37 °C, and induced with 1 mM IPTG. Induced cells were incubated at 16 °C for 16 h with shaking and then harvested by centrifugation at 4,000 rpm. Cells were lysed by two passes in a LM20 Microfluidizer at 18,000 psi in lysis buffer (50 mM Tris-HCl pH 7.4, 200 mM NaCl, 10 mM MgCl2) supplemented with 0.5 mg/mL lysozyme and 1 mM PMSF. Clarified lysate was incubated with Glutathione Superflow Agarose (Pierce) to bind the GST tag. Protein bound to the beads was eluted with the addition of 10 mM reduced glutathione. Eluate was further purified on a 1 mL HiTrap Q IEX column. Peak fractions were pooled, concentrated, and then supplemented with 10% glycerol before snap freezing.

The list of constructs used in this study and their denatured PAGE gels after purification can be found in **Extended Data Table 1** and **Extended Data Fig. 9**, respectively.

## Protein Labeling

Porcine brain tubulin was labeled with NHS reactive dyes or fluorophores, as previously described^71^. MTs were polymerized in BRB80 supplemented with 1 mM GTP and 1 mM DTT for 30 min at 37 °C, then pelleted over a high pH glycerol cushion (100 mM HEPES pH 8.6 1 mM MgCl2, 1 mM EGTA, 1 mM GTP, 1 mM DTT, 60 % glycerol) at 80,000 rpm at 37 °C in a TLA100.4 rotor. The MT pellet was washed and resuspended in labeling buffer (100 mM HEPES pH 8.6, 1 mM MgCl_2_, 1 mM EGTA, 1 mM GTP, 1 mM DTT, 40 % glycerol) and incubated with 5x molar excess NHS-label at 37 C for 30 min. Labeled MTs were pelleted over a low pH cushion (BRB80 supplemented with 1 mM GTP, 1 mM DTT, 60 % glycerol) at 80,000 rpm at 37 °C in a TLA100.4 rotor. The labeled MT pellet was washed with warm BRB80, then resuspended in ice-cold BRB80 and incubated on ice for 15 min. The final protein concentration and labeling efficiency were quantified using a Nanodrop spectrophotometer before snap freezing in LN2.

MAPs with a ybbR tag were labeled, as previously described^45^. Briefly, protein was incubated with 2x molar excess of CoA-derivatized dyes for 1 h at room temperature in 200 µL SFP buffer supplemented with 0.2 mg/mL SFP enzyme. Proteins with a SNAP-tag were labeled by incubation with 5x molar excess of benzylguanine-derivatized dye at room temperature for 1 h.

## MT Polymerization

Prior to all MT polymerizations, tubulin was thawed from -80 °C and clarified by a 10 min spin at 100,000 rpm in a TLA 100 rotor at 4 °C. Taxol-stabilized MTs were polymerized by stepwise Taxol addition. 5% biotin- or dye-labeled porcine tubulin was added to the polymerizing tubulin for immobilizing and visualizing MTs in TIRF microscopy. Tubulins were incubated with BRB80 supplemented with 1 mM GTP and DTT on ice before polymerization. Tubulin was then moved to a 37 °C water bath, and 0.1 μM Taxol was added to the solution. Polymerizing MTs were incubated for 10 min before the addition of 1 μM Taxol, then another 10 min before the addition of 10 μM Taxol. After 15 min, polymerized MTs were pelleted at 20,000 rcf and resuspended in BDT (BRB80 supplemented with 1 mM DTT and 10 μM Taxol). Porcine brain MTs were kept in the dark for up to two weeks at room temperature. MTs made from recombinant tubulin were used within two days of polymerization. For CryoEM, MTs were polymerized without labeled tubulin. PelA-stabilized MTs were prepared using stepwise addition of 0.01, 0.1, and 1 μM PelA with the same time and temperature intervals as Taxol.

GMPCPP-stabilized MT seeds were prepared by incubating 1 mg/mL tubulin in BRB80 supplemented with 1 mM DTT and 1 mM GMPCPP for 10 min on ice. Afterwards, 5% DMSO was added to the solution, and the MTs were polymerized for 45 min by incubation in a 37 °C water bath. Polymerized MTs were pelleted at 20,000 rcf and resuspended in BRB80 + 1 mM DTT.

For kinesin-1 motility assays, longer Taxol-stabilized MTs were polymerized on functionalized coverslips. Biotinylated GMPCPP MT seeds were immobilized to biotin-PEG coverslips via streptavidin. Then, 1.6 mg/mL tubulin in polymerization buffer (PB) (BRB80 supplemented with 0.2% methylcellulose, 1mg/mL casein, 0.5% Pluronic F-127, 1 mM GTP, 1 mM DTT, 1% glucose oxidase, catalase, 0.4% dextrose) was washed over the seeds and warmed to 37 °C for 10 min. The polymerized MTs were then washed with warm PB lacking tubulin and supplemented with 10 μM Taxol and incubated at 37 °C for 10 min before imaging.

## In vitro Glutamylation

Taxol-stabilized α1A/β3 MTs were incubated overnight in glutamylation buffer (20 mM HEPES pH 7.0, 50 mM NaCl, 5 mM MgCl_2_, 1 mM ATP, 1 mM glutamate, 1 mM DTT, 10 μM Taxol) with purified TTLL7 at a 1:10 molar ratio (TTLL7:tubulin). After incubation, MTs were pelleted and resuspended in BRB80 supplemented with 350 mM NaCl, 1 mM GTP, 1 mM DTT, 10 μM Taxol and incubated at 37 °C for 15 min to unbind TTLL7. MTs were pelleted again and resuspended in cold BRB80 on ice for 15 min to depolymerize MTs into tubulin. The glutamylated tubulin solution was supplemented with 10% glycerol and snap frozen.

## α1A/β3 in vitro MT Characterization

Taxol-stabilized MTs were diluted twenty-fold into BDT and immobilized via streptavidin onto PEG-biotin surfaces. Bundled MTs were determined as either having greater than three times the average single-MT intensity or by having an intersection of 3 or more MTs. MT stability was assayed by washing out Taxol and imaging over 10 min to watch the degradation of MTs. MT length was determined 24h after polymerization.

## SiR-Tubulin Binding

Taxol-stabilized MTs polymerized from α1A/β3 or porcine brain tubulin were co-polymerized with tubulin labeled with Cy3 and Atto488, respectively. Both MTs were immobilized to the coverslip via streptavidin in BRB80 supplemented with 10 μM Taxol and 1 mM DTT. The Taxol was washed out and replaced with 200 nM SiR-tubulin. SiR-tubulin intensity colocalizing to different MT labels was quantified and compared with a two-sample t-test.

## MAP-MT Binding Affinity

To quantify the binding of MAPs to stabilized MTs, Taxol- or PelA-stabilized MTs were immobilized to PEG-biotin coverslips using streptavidin. Increasing concentrations of labeled MAP protein were flown into the chamber in imaging buffer (IB; 30 mM HEPES pH 7.4, 5 mM MgSO_4_, 1 mM EGTA, 0.1% MCell, casein, 0.5% Pluronic F-127, 5 mM TCEP, glucose oxidase, catalase, 0.4% dextrose, 150 mM KAc) supplemented with 10 μM Taxol or 1 μM PelA and the fluorescence intensity of the MAP colocalizing to the MTs was measured. The MTIMBS software (available at Yildiz-Lab GitHub) was used to segment images, measure the average fluorescence intensity of MTs, and subtract local background. Fluorescent intensities were plotted against concentration and fitted to a Hill equation with n = 1 for comparison.

To quantify MT binding of MAPs on dynamically growing MTs, GMPCPP seeds were immobilized to the coverslip and 1.6 mg/mL porcine brain tubulin, along with 100 nM of tau or MAP7 in PB, were added to the chamber. Chambers were sealed with nail polish and warmed to 37 °C. After 10 min of MT growth, images were acquired to quantify the intensities of individual MT extensions and seeds, and the ratio of seed intensity to extension intensity for each MT.

To determine the binding of tau to GDP MTs without stabilizing agents or tubulin in solution, MTs were grown from GMPCPP seeds in PB, as described above. After 15 min of growth, the tubulin-containing solution was washed out of the chamber and replaced with 100 nM tau in PB. Tau intensities were quantified on the third frame after flow (15 s later) and compared via a two-sample t-test.

## Kinesin-1 Motility Assays

Long Taxol-stabilized MTs were prepared in imaging chambers, as described above. 2 nM LD655-labeled (Lumidyne) K560-GFP-SNAP was added to the chamber in IB supplemented with 1 mM ATP. Kinesin-1 run frequencies were determined by the number of processive kinesin-1 motors that landed per length of MT per unit time. As an internal control, kinesin-1 run frequencies were normalized to the run frequency of kinesins that landed on the GMPCPP seeds in each experiment.

## Two MAP Competition Assays

Stabilized MTs were immobilized to PEG-biotin coverslips via biotin-streptavidin linkage in three separate chambers, containing either MAP or both in 500 nM concentration. The intensity of each MAP in the competitive chambers is compared to the intensity in the control chambers and reported as a percent recovery. For the MAP9 vs MAP4 and tau vs MAP7 competitions in the presence of PelA, 100 nM of each MAP was used.

## MAP/Motor Motility Assays

Long Taxol-stabilized MTs were grown from GMPCPP seeds using either α1A/β3 or porcine brain tubulin. 2 nM LD655-labeled K560-GFP-SNAP was added to the chamber in IB supplemented with 1 mM ATP with equimolar concentrations of MAP7 and tau in solution. Kinesin-1 run frequencies were determined by the number of kinesin-1 molecules that landed per length of MT per unit time.

## Tau Washout Assay

To determine tau dwell time on compacted lattices, PelA-MTs were immobilized to a coverslip via streptavidin and incubated with 100 nM tau in IB for 5 min. Before imaging, the chamber was washed with IB lacking tau. MTs were imaged every 5 s at 200 ms exposure for 7 min.

## Tau Diffusion Assay

GMPCPP or PelA-stabilized MTs were immobilized on a coverslip via streptavidin. GMPCPP MTs were incubated with 10 nM SNAP-tau labeled with LD655 in IB. PelA MTs were incubated with 15 pM SNAP-tau labeled with LD655 in IB. Images were collected constantly for 30 s at 200 ms exposure time.

## Tau Envelope Growth Assay

To assay tau variants’ capacity to compact the MT lattice, GMPCPP- or Taxol-stabilized MTs were immobilized to coverslips. Short extensions of compacted GDP-lattice MTs were grown from the seeds by incubating the coverslips with 1.6 mg/mL α1A/β3 tubulin in PB at 37 °C for 1 min, followed by washout of free tubulin using PB supplemented with 10 μM Taxol. The chamber was incubated for 5 min at 37 °C and then loaded with 500 nM tau in IB and imaged immediately to watch condensate growth from the compacted MT extensions.

## MT Curvature and MAP Intensity Measurements

From MAP competition assays containing 100 nM tau and 100 nM MAP7 in IB, curved MTs were segmented using MTIMBS. The x-y positions of each pixel were fitted to a 7th-degree polynomial, and the curvature at each pixel was calculated using the first and second derivatives. Average fluorescence intensity over background in each color channel was measured per MT, then each pixel intensity was normalized by dividing by the average intensity in that channel. Pixel intensities were then plotted for each channel as a function of curvature.

## Cryo-EM Sample Preparation

Recombinant naive microtubules were polymerized through the same methods mentioned above, stabilized with taxol, and diluted to 5μM. 4 μL of 5 μM taxol-recombinant MTs were incubated on a glow discharged holey carbon cryo-EM grid (QuantiFoil, Cu 300 R 1.2/1.3) inside the chamber of the Vitrobot for 30 s, set at 25 °C and 80% humidity, plunge-frozen in liquid ethane with a blot force of 5 units and a blot time of 6 s, and transferred to liquid nitrogen.

## Cryo-EM Data Collection

Cryo-EM data were collected on an Arctica microscope (Thermo Fisher Scientific) operated at 200 kV with a K3 direct electron detector (Gatan). Images were acquired at 36,000x magnification at 1.14 Å/pixel and were acquired in super-resolution mode with a dose rate of ∼7.2 electrons/pixel/second and exposure time of ∼9 s dose-fractionated into 50 frames. All data were collected using the SerialEM software package^72^.

## Cryo-EM Image Processing

Data processing followed established protocols for MT cryo-EM^73, 74^. Movie stacks were motion-corrected in cryoSPARC^75^ and CTF parameters were estimated with the patch CTF job. Micrographs with poor CTF fits were excluded manually. Particles were picked automatically with the filament tracer, initially without a template and then using 2D class averages, with a segment separation of 82 Å. Particle images were extracted at 512 px and subjected to four rounds of 2D classification. Heterogeneous refinement against 13- and 14-protofilament MT references was used to separate the two populations. The particles were approximately evenly between the two classes, and the 14-protofilament class was used for all downstream processing. 14-protofilament particles were subjected to helical refinement (initial rise 82.5 Å, twist 0°) followed by local refinement with a cylindrical MT mask. Helical parameters were refined with the symmetry search job and used for pseudo-helical symmetry expansion. Local refinement on the expanded particle stack yielded the symmetrized MT reconstruction. Focus 3D classification without alignment of the reconstruction using three classes revealed two classes with evenly distributed particle number and sorted distribution of alpha and beta tubulin. One class was arbitrarily selected and underwent a volume align shift job in cryoSPARC, shifting the particle stack by the length of approximately one tubulin monomer in real space (41.5 Å). The two classes were therefore aligned in distributions of alpha and beta tubulins and were combined in a local 3D refinement. The resulting reconstruction was a properly sorted 3D reconstruction of the recombinant microtubule at 3.67 Å resolution.

---

**Extended Data Fig. 1.**
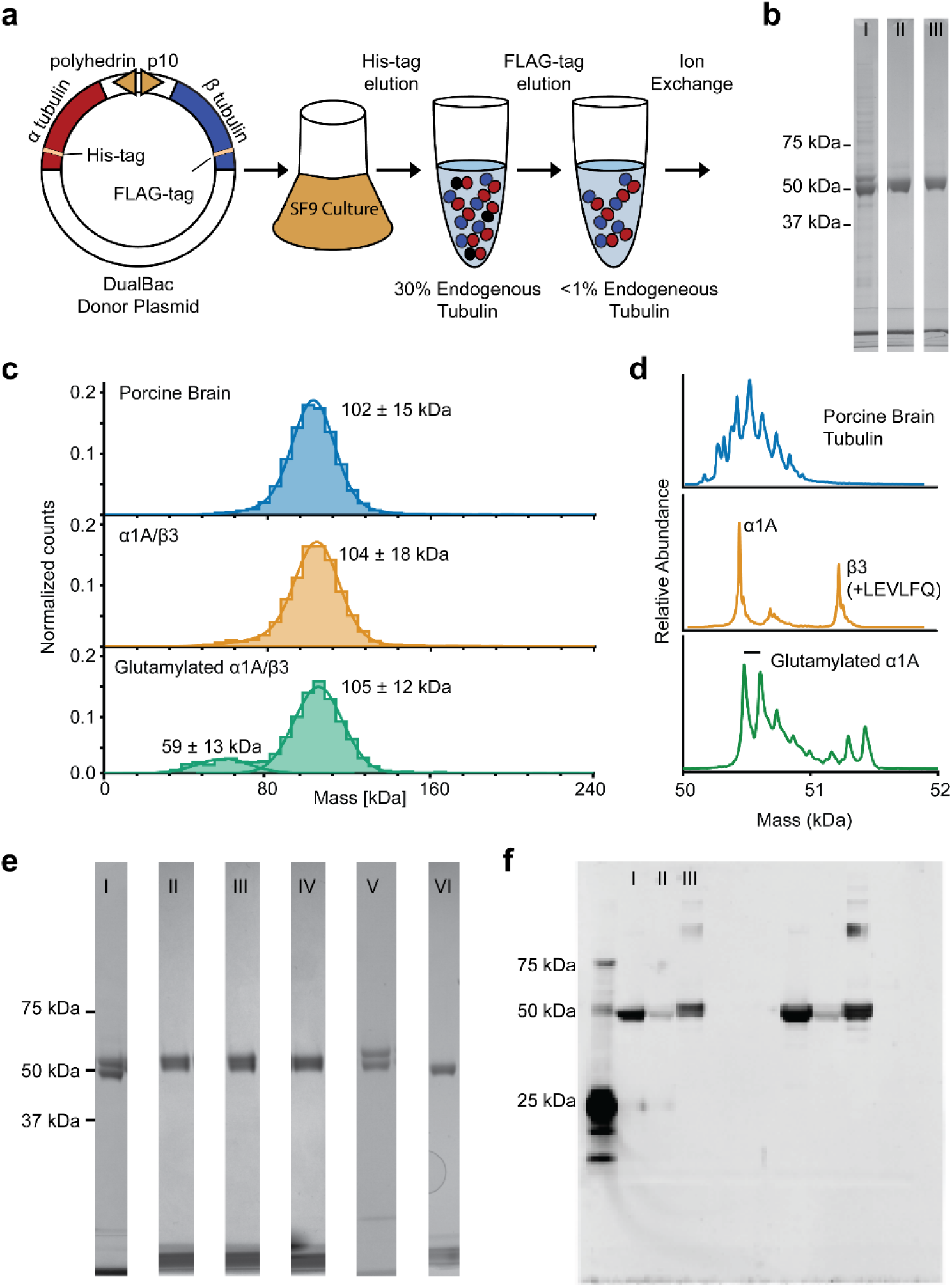
**Expression and purification of recombinant tubulin**. **a,** Purification scheme for α1A/β3 tubulin. **b,** Representative gel images of α1A/β3 tubulin purification after I) His-tag, II) FLAG-tag, and III) ion exchange purification steps. **c,** Mass photometry of purified tubulin proteins. **d,** Mass spectra of porcine brain tubulin (top), α1A/β3 tubulin (middle), and in vitro glutamylated α1A/β3 tubulin (bottom). **e,** Representative gel images of I) α tailless, II) naïve, III) detyrosinated, IV) delta two, V) lumen-tagged α1A/β3, and VI) porcine brain tubulin. **f,** Western blot using gt335 antibody against glutamylated tubulin for I) porcine brain, II) α1A/β3, and III) in vitro glutamylated α1A/β3 tubulin.

**Extended Data Fig. 2.**
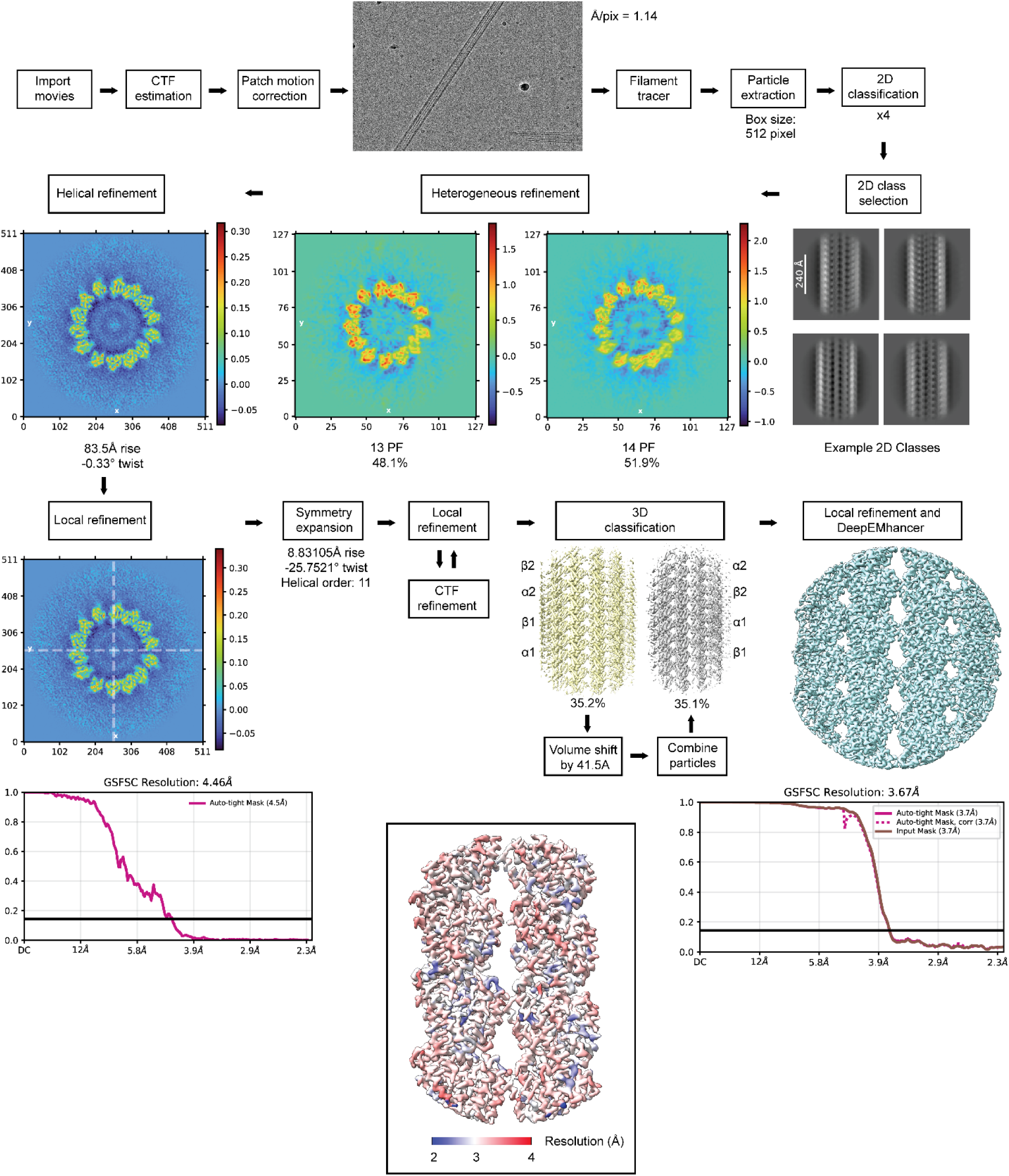
Cryo-EM processing pipeline for naive recombinant α1A/β3 microtubule structure. The analysis pipeline reveals the local resolution map and the FSC plot for the recombinant microtubule structure.

**Extended Data Fig. 3.**
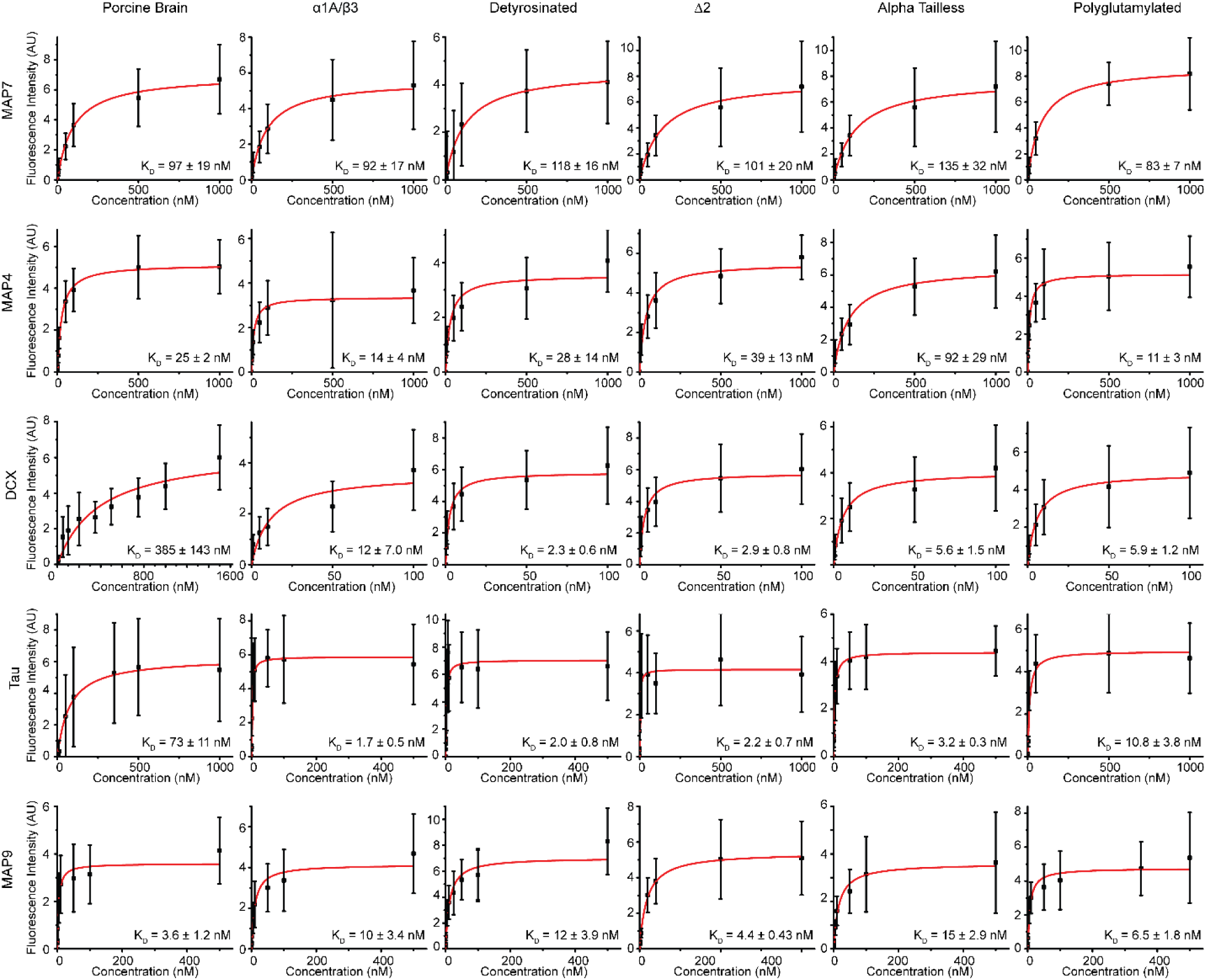
**Affinity of structural MAPs to MTs polymerized with recombinant or porcine brain tubulin**. Taxol-MTs polymerized from tubulins as denoted from left to right: porcine brain, α1A/β3, detyrosinated, delta two, α tailless, and in vitro glutamylated tubulin. MAP protein binding from different MAPs from top to bottom: MAP7, MAP4, DCX, tau, and MAP9. Black squares denote mean fluorescence intensity from 200 MTs across two technical replicates, error bars denote SD. Reported K_D_ values are the best fit to a Hill function with n=1, ± SE.

**Extended Data Fig. 4.**
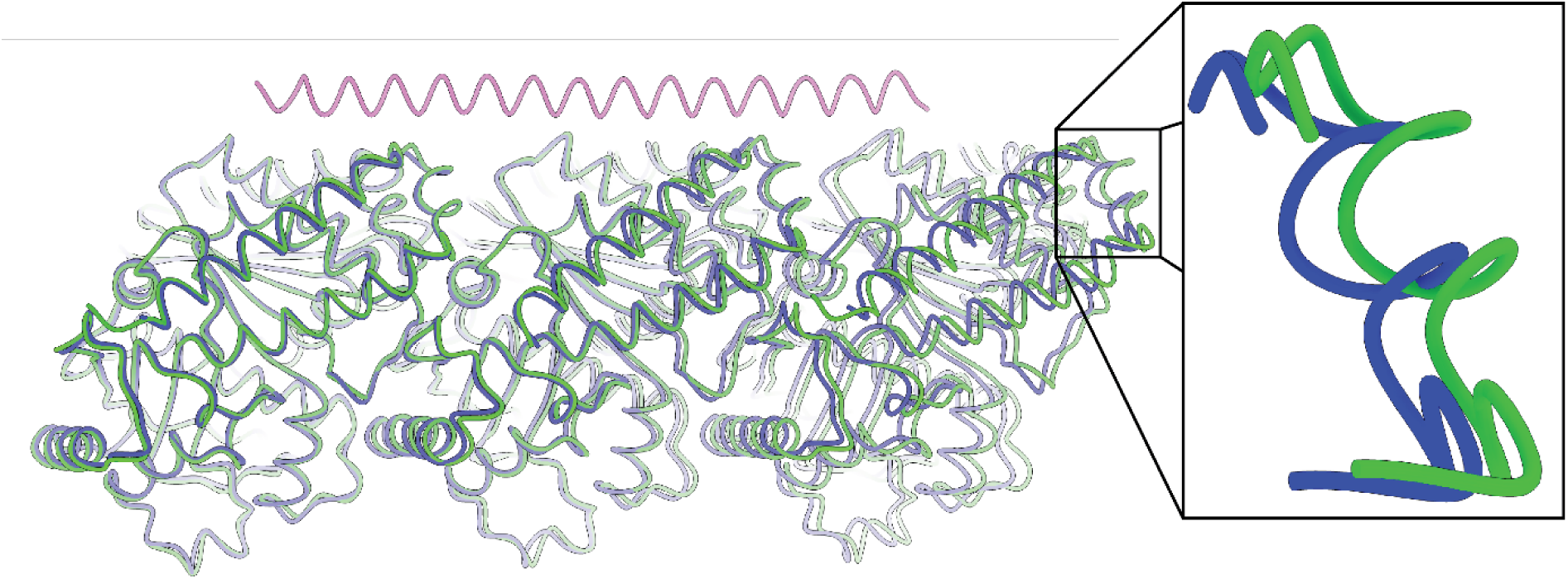
MAP7 expands the MT lattice. Published structures of undecorated PelA-stabilized MTs (blue, PDB# 5SYC^47^) versus MAP7 (pink) bound PelA-stabilized MTs (green, PDB# 7SGS^45^) aligned by the minus-end alpha tubulin. Insert: The second alpha tubulin H11 is translated by approximately 1 Å away, indicating an expanded MT lattice.

**Extended Data Fig. 5.**
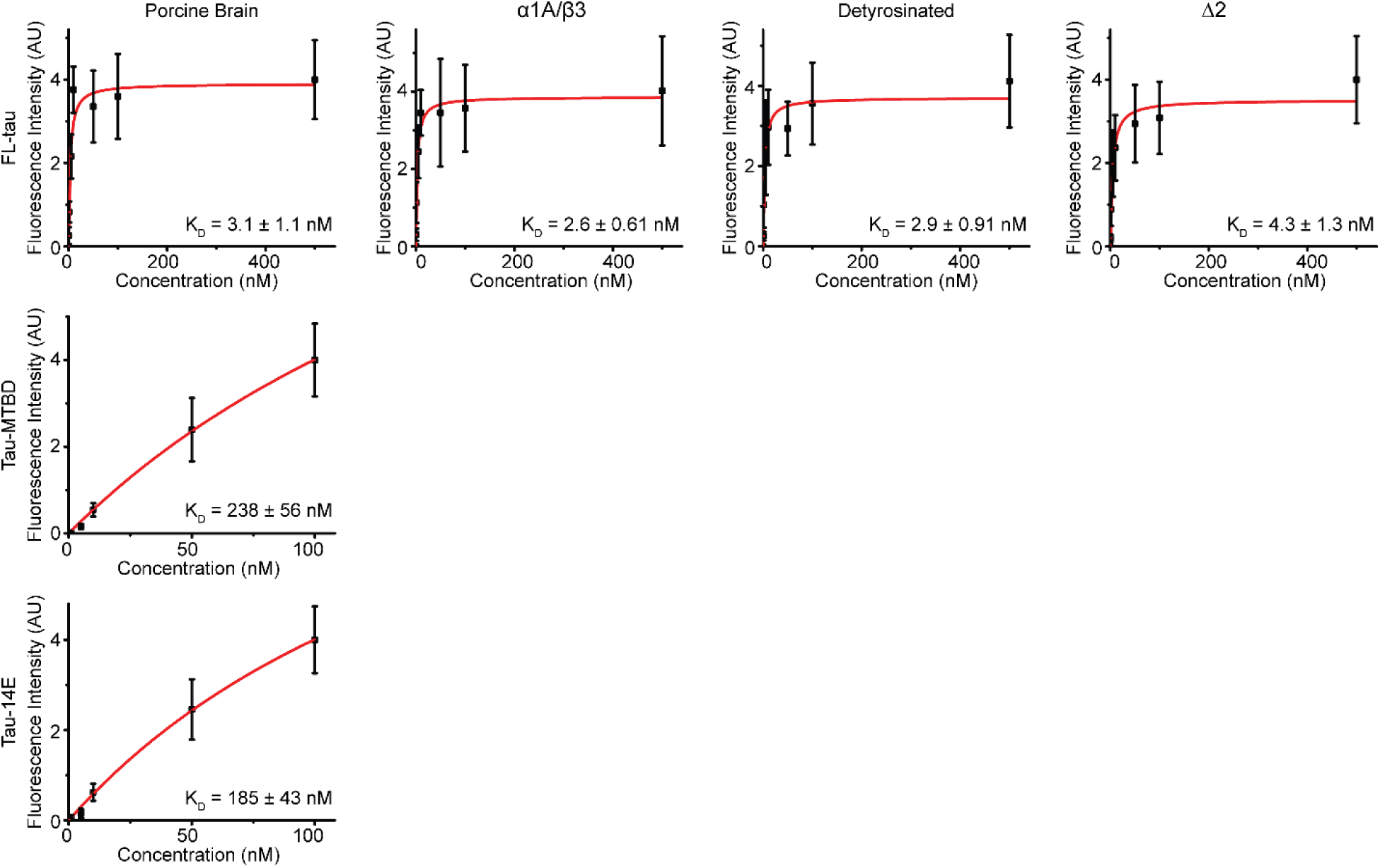
**Tau binding to PelA-stabilized MTs**. PelA-MTs polymerized from tubulins as denoted from left to right: porcine brain, α1A/β3, detyrosinated, and Δ2 tubulin. Tau variants as denoted from top to bottom: FL tau, tau-MTBD, and tau-14E. Black squares denote mean fluorescence intensity from 50 MTs across two technical replicates; error bars denote SD. Reported K_D_ values are the best fit to a Hill function with n=1, ± SE.

**Extended Data Fig. 6.**
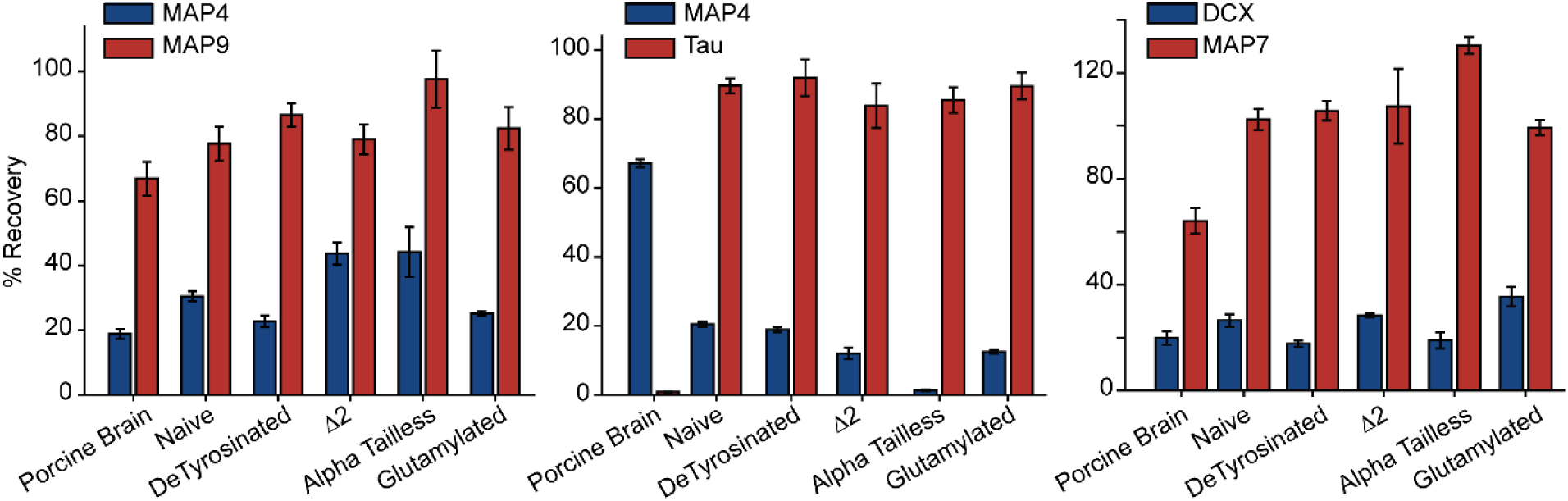
**Two MAP competition assays**. The MT binding competition assay between MAP4 vs MAP9, MAP4 vs tau, and DCX vs MAP7. Bars represent mean ± SD from three technical replicates.

**Extended Data Fig. 7.**
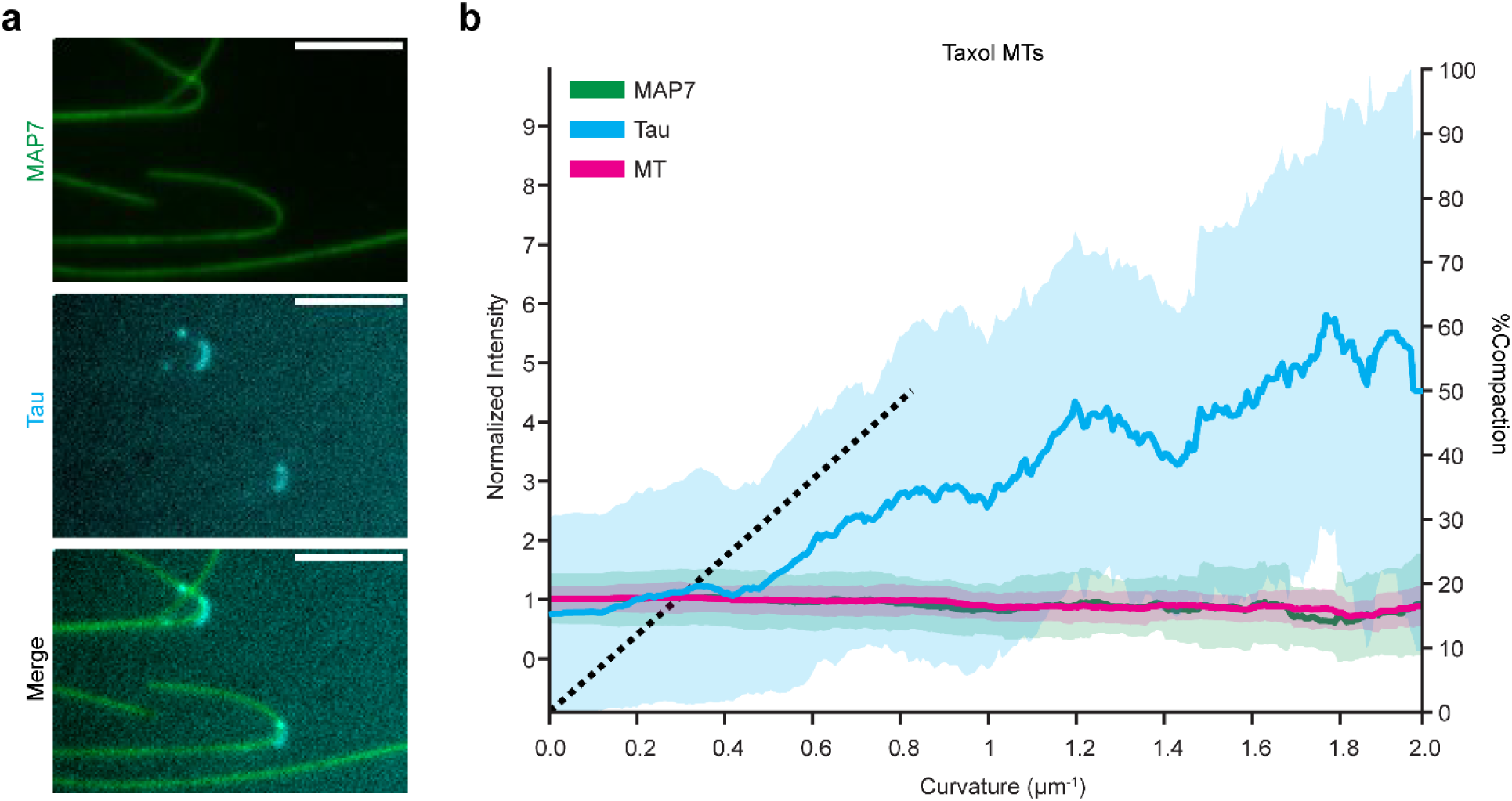
MAP7 vs Tau competition on Taxol-stabilized MTs. a, Representative images of 100 nM tau (cyan) and MAP7 (green) on MTs, scale bars: 5 μm. **b,** Fluorescence intensity as a function of MT curvature for MAP7 and tau on Taxol MTs. MAP7 in green, tau in cyan, MT in magenta. Thick bars represent mean fold-increase over average MT intensity, and shaded regions denote standard error. Data from 84 curved MTs across two technical replicates.

**Extended Data Fig. 8.**
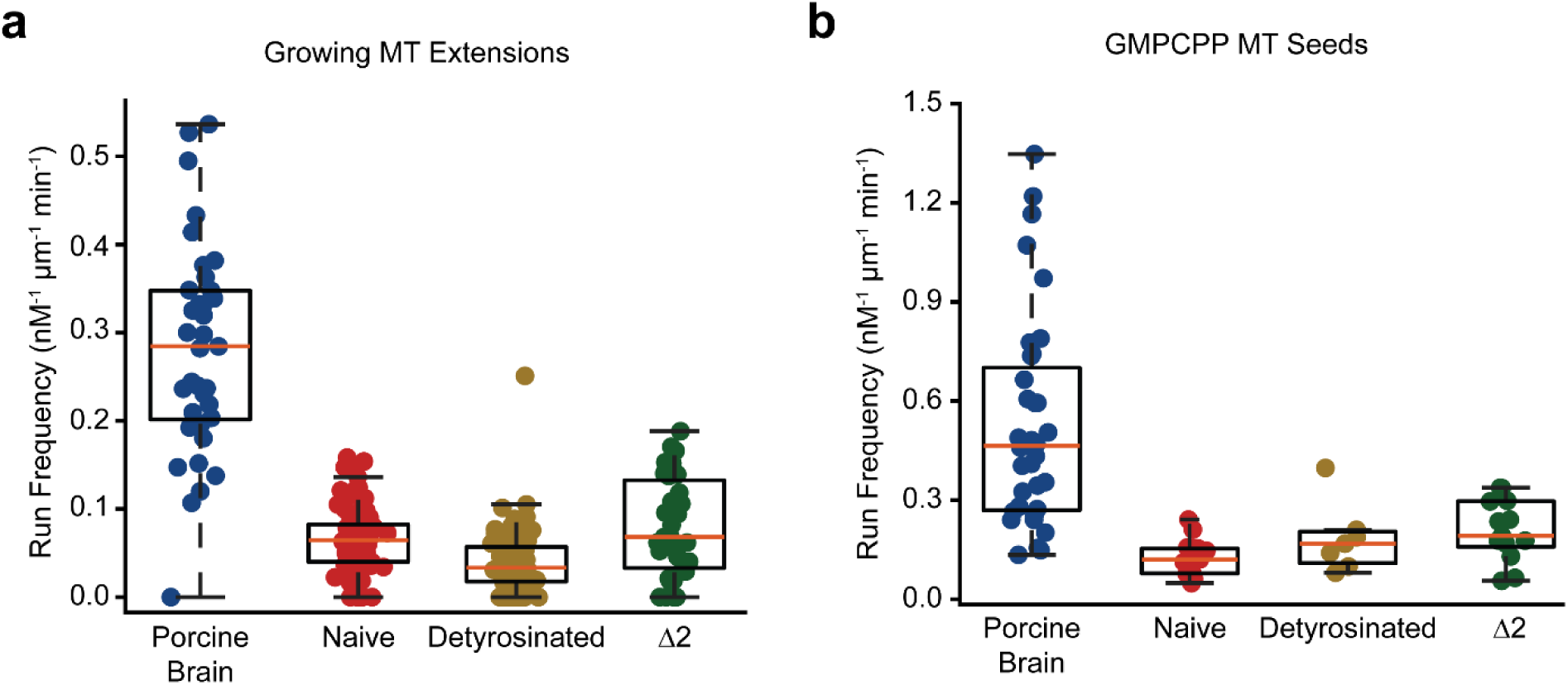
Tubulin in solution impacts kinesin-1 run frequency. a, Run frequency of single kinesin-1 motors on the dynamically growing MT extensions made from different tubulins in solution. From left to right, N = 35, 81, 100, 35 MTs across two technical replicates. **b,** Run frequency of single kinesin-1 motors on porcine brain GMPCPP-stabilized MT seeds longer than 5 μm, with different tubulins in solution. From left to right, N = 36, 18, 7, 13 MT seeds across two technical replicates.

**Extended Data Fig. 9.**
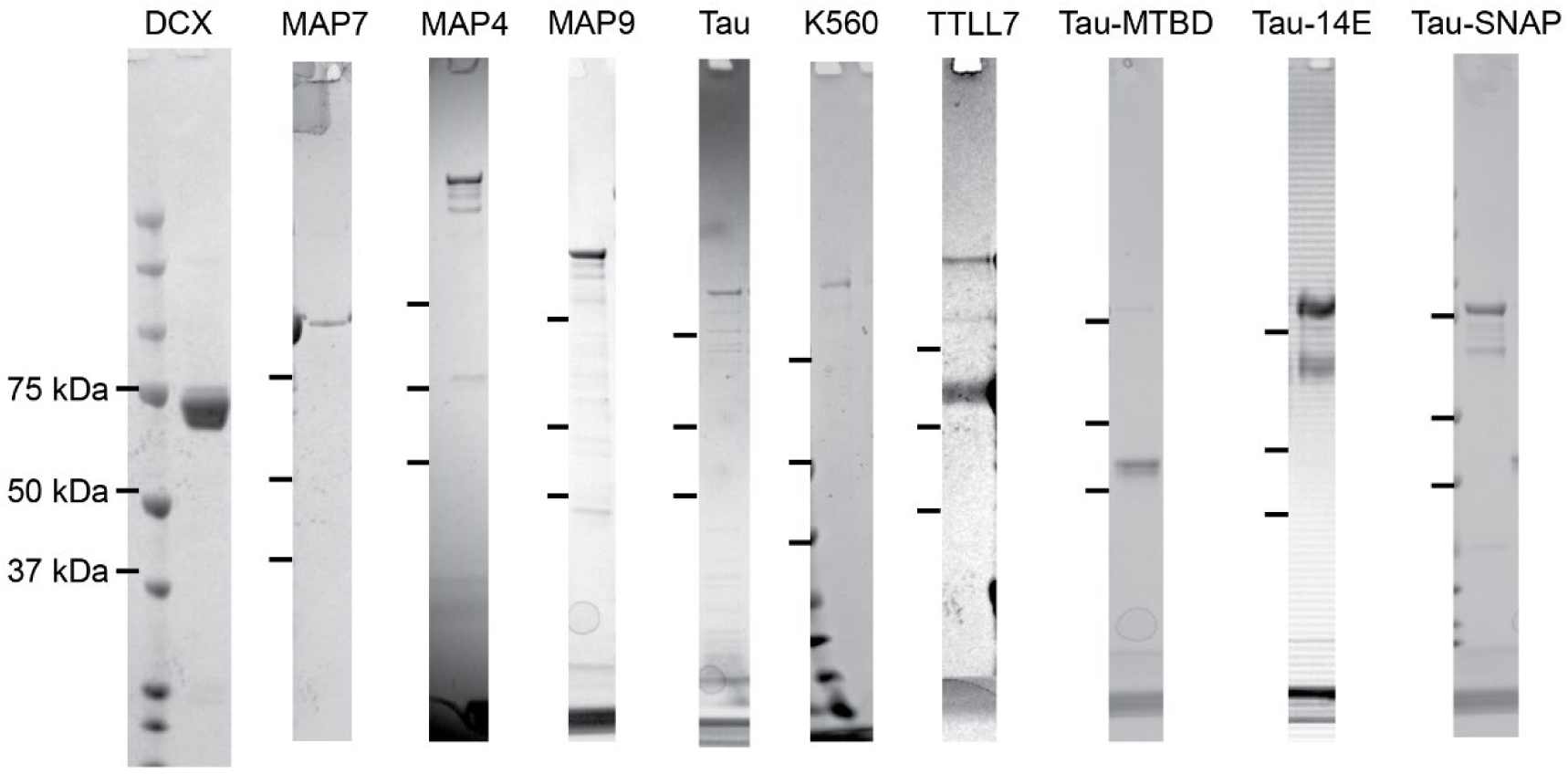
**Denatured gel pictures of purified proteins**. PAGE gel images of purified MAPs, kinesin, and TTLL7 used in this study.

**Extended Data Table 1.** The list of constructs used in this study. Amino acid numbers and point mutations are shown in subscripts and superscripts, respectively.

| Name | Description | Source | Expression | Figures |
| --- | --- | --- | --- | --- |
| α1A/β3 Tubulin | α1ATUB<sup>1-36</sup>:: 8xHis:: α1ATUB<sup>45-452</sup><br>β3TUB<sup>1-450</sup>:: HRV 3C:: FLAG | 70 | SF9 | 1B-G, 2C-F,H, 4B,D,F, 5B,D,E, S1C-F, S2, S3, S4, S5 |
| Lumen-tagged α1A/β3 | α1ATUB<sup>1-36</sup>:: 8xHis:: α1ATUB<sup>45-452</sup><br>β3TUB<sup>1-38</sup>:: GGGS:: FLAG:: GGGGSG:: β3TUB<sup>39-450</sup> | This study | SF9 | 1B-H |
| Detyrosinated α1A/β3 | α1ATUB<sup>1-36</sup>:: 8xHis:: α1ATUB<sup>45-451</sup><br>β3TUB<sup>1-450</sup>:: HRV 3C:: FLAG | 70 | SF9 | 2C,D,F, 4B,D, 5B, S1B, S2, S3, S4, S5 |
| Delta Two α1A/β3 | α1ATUB<sup>1-36</sup>:: 8xHis:: α1ATUB<sup>45-450</sup><br>β3TUB<sup>1-450</sup>:: HRV 3C:: FLAG | 70 | SF9 | 2C,D,F, 4B,D, 5B, S2, S3, S4, S5 |
| Alpha Tailless α1A/β3 | α1ATUB<sup>1-36</sup>:: 8xHis:: α1ATUB<sup>45-439</sup><br>β3TUB<sup>1-450</sup>:: HRV 3C:: FLAG | 70 | SF9 | 2C,D, 5B, S2, S5 |
| Kinesin-1 | ZZ:: TEV:: KIF5B<sup>1-560</sup> :: GFP:: SNAP | 45 | SF9 | 4A-F, 5D-G, S4 |
| MAP7 | ZZ:: TEV:: ybbR:: MAP7<sup>1-749</sup> | 45 | SF9 | 1F, 2A,D,G, 5D-G, S2, S5 |
| MAP7-mStayGold | ZZ:: TEV:: ybbR:: MAP7<sup>1-749</sup>:: mStayGold | This study | SF9 | 5C |
| 2N4R Tau | 6xHis:: TwinStrepII:: sfGFP:: 2N4R Tau | 62 | Rosetta2 DE3 | 1F, 2D-H, 3A-C,E,H, 5D-G, S2, S3, S5 |
| Tau-MTBD | 6xHis:: TwinStrepII:: sfGFP:: 2N4R Tau<sup>244-373</sup> | This study | Rosetta2 DE3 | 3A-C, S3 |
| Tau-14E | 6xHis:: TwinStrepII:: sfGFP:: 2N4R Tau<sub>T111E, T153E, T175E, T181E, S199E, S202E, T205E, T212E, T217E, T231E, S235E, S396E, S404E, S422E</sub> | This study | Rosetta2 DE3 | 3A-C, F, S3 |
| SNAP-Tau | 6xHis:: TwinStrepII:: SNAP:: 2N4R Tau | This study | Rosetta2 DE3 | 3G, 5C |
| DCX | 6xHis:: HRV 3C:: DCX:: eGFP:: StrepTag | 38 | Rosetta2 DE3 | 2D, S2, S5 |
| MAP4 | 6xHis:: TwinStrepII:: SNAP:: HRV 3C:: MAP4:: FLAG | 38 | SF9 | S2, S5 |
| MAP9 | 6xHis:: TwinStrepII:: sfGFP:: HRV 3C:: MAP9:: FLAG | 38 | Rosetta2 DE3 | S2, S5 |
| TTLL7 | GST:: TEV:: TTLL7<sup>1-518</sup> | 54 | Rosetta2 DE3 | S1C,D,F |

## References

1. Barlan, K. & Gelfand, V.I. Microtubule-Based Transport and the Distribution, Tethering, and Organization of Organelles. LID - 10.1101/cshperspect.a025817 [doi] LID - a025817.

2. Gross, S.P. Hither and yon: a review of bi-directional microtubule-based transport.

3. Janke, C. & Magiera, M.M. The tubulin code and its role in controlling microtubule properties and functions. Nature reviews. Molecular cell biology 21, 307–326 (2020).

4. Verhey, K.J. & Gaertig, J. The tubulin code. Cell cycle 6, 2152–2160 (2007).

5. Hausrat, T.J., Radwitz, J., Lombino, F.L., Breiden, P. & Kneussel, M. Alpha- and beta- tubulin isotypes are differentially expressed during brain development. Dev Neurobiol 81, 333–350 (2021).

6. Luduena, R.F. Multiple forms of tubulin: Different gene products and covalent modifications. Int Rev Cytol 178, 207–275 (1998).

7. Garnham, C.P. et al. Multivalent Microtubule Recognition by Tubulin Tyrosine Ligase-like Family Glutamylases. Cell 161, 1112–1123 (2015).

8. Janke, C. et al. Tubulin polyglutamylase enzymes are members of the TTL domain protein family. Science 308, 1758–1762 (2005).

9. Ti, S.-C., Alushin, G.M. & Kapoor, T.M. Human β-Tubulin Isotypes Can Regulate Microtubule Protofilament Number and Stability. Dev Cell 47, 175–190.e175 (2018).

10. Schwer, H.D. et al. A lineage-restricted and divergent &#x3b2;-tubulin isoform is essential for the biogenesis, structure and function of blood platelets. Curr Biol 11, 579–586 (2001).

11. Vemu, A. et al. Structure and Dynamics of Single-isoform Recombinant Neuronal Human Tubulin. The Journal of biological chemistry 291, 12907–12915 (2016).

12. Portran, D., Schaedel, L., Xu, Z., Théry, M. & Nachury, Maxence V. Tubulin acetylation protects long-lived microtubules against mechanical ageing. Nat Cell Biol 19, 391–398 (2017).

13. Xu, Z. et al. Microtubules acquire resistance from mechanical breakage through intralumenal acetylation. Science 356, 328–332 (2017).

14. McKenney, R.J., Huynh, W., Vale, R.D. & Sirajuddin, M. Tyrosination of alpha-tubulin controls the initiation of processive dynein-dynactin motility. The EMBO journal 35, 1175–1185 (2016).

15. Fan, X. & McKenney, R.J. Control of motor landing and processivity by the CAP-Gly domain in the KIF13B tail. Nature communications 14, 4715 (2023).

16. Valenstein, M.L. & Roll-Mecak, A. Graded Control of Microtubule Severing by Tubulin Glutamylation. Cell 164, 911–921 (2016).

17. Genova, M. et al. Tubulin polyglutamylation differentially regulates microtubule-interacting proteins. Embo J 42 (2023).

18. Gadadhar, S. et al. Tubulin glycylation controls axonemal dynein activity, flagellar beat, and male fertility. Science 371, 144-+ (2021).

19. Reed, N.A. et al. Microtubule acetylation promotes kinesin-1 binding and transport. Current biology : CB 16, 2166–2172 (2006).

20. Konishi, Y. & Setou, M. Tubulin tyrosination navigates the kinesin-1 motor domain to axons. Nature neuroscience 12, 559–567 (2009).

21. Tas, R.P. et al. Differentiation between Oppositely Oriented Microtubules Controls Polarized Neuronal Transport. Neuron 96, 1264–1271 e1265 (2017).

22. Thorn, K.S., Ubersax, J.A. & Vale, R.D. Engineering the processive run length of the kinesin motor. J Cell Biol 151, 1093–1100 (2000).

23. Wang, Z. & Sheetz, M.P. The C-terminus of tubulin increases cytoplasmic dynein and kinesin processivity. Biophysical journal 78, 1955–1964 (2000).

24. Serrano, L., Avila, J. & Maccioni, R.B. Controlled Proteolysis of Tubulin by Subtilisin - Localization of the Site for Map2 Interaction. Biochemistry-Us 23, 4675–4681 (1984).

25. Sirajuddin, M., Rice, L.M. & Vale, R.D. Regulation of microtubule motors by tubulin isotypes and post-translational modifications. Nature cell biology 16, 335–344 (2014).

26. Lakämper, S. & Meyhöfer, E. The E-hook of tubulin interacts with kinesin’s head to increase processivity and speed.

27. Shigematsu, H. et al. Structural insight into microtubule stabilization and kinesin inhibition by Tau family MAPs. J Cell Biol 217, 4155–4163 (2018).

28. Paquette, A.L. et al. Competition for microtubule lattice spacing between a microtubule expander and compactor. Curr Biol 35, 4442–4452.e4444 (2025).

29. Alushin, G.M. et al. High-Resolution Microtubule Structures Reveal the Structural Transitions in αβ-Tubulin upon GTP Hydrolysis. Cell 157, 1117–1129 (2014).

30. Vangos, N., et al. Taxol exploits molecular switches within tubulin to stabilize microtubules. *bioRxiv* (2026).

31. Zhang, R., Alushin, G.M., Brown, A. & Nogales, E. Mechanistic Origin of Microtubule Dynamic Instability and Its Modulation by EB Proteins. Cell 162, 849–859 (2015).

32. LaFrance, B.J. et al. Structural transitions in the GTP cap visualized by cryo-electron microscopy of catalytically inactive microtubules. Proceedings of the National Academy of Sciences of the United States of America 119 (2022).

33. Shen, Y.S. & Ori-McKenney, K.M. Microtubule-associated protein MAP7 promotes tubulin posttranslational modifications and cargo transport to enable osmotic adaptation. Dev Cell 59 (2024).

34. Manka, S.A.-O.X. & Moores, C.A.-O. The role of tubulin-tubulin lattice contacts in the mechanism of microtubule dynamic instability.

35. Siahaan, V. et al. Microtubule lattice spacing governs cohesive envelope formation of tau family proteins. Nat Chem Biol 18, 1224–1235 (2022).

36. Nakata, T., Niwa, S., Okada, Y., Perez, F. & Hirokawa, N. Preferential binding of a kinesin- 1 motor to GTP-tubulin-rich microtubules underlies polarized vesicle transport. J Cell Biol 194, 245–255 (2011).

37. Shima, T. et al. Kinesin-binding-triggered conformation switching of microtubules contributes to polarized transport. J Cell Biol 217, 4164–4183 (2018).

38. Monroy, B.Y. et al. A Combinatorial MAP Code Dictates Polarized Microtubule Transport. Developmental cell 53, 60–72 e64 (2020).

39. Ebneth, A. et al. Overexpression of tau protein inhibits kinesin-dependent trafficking of vesicles, mitochondria, and endoplasmic reticulum: implications for Alzheimer’s disease. J Cell Biol 143, 777–794 (1998).

40. Gumy, L.F. et al. MAP2 Defines a Pre-axonal Filtering Zone to Regulate KIF1- versus KIF5-Dependent Cargo Transport in Sensory Neurons. Neuron 94, 347–362 e347 (2017).

41. Semenova, I. et al. Regulation of microtubule-based transport by MAP4. Mol Biol Cell 25, 3119–3132 (2014).

42. Siahaan, V. et al. Kinetically distinct phases of tau on microtubules regulate kinesin motors and severing enzymes. Nature cell biology 21, 1086–1092 (2019).

43. Chaudhary, A.R. et al. MAP7 regulates organelle transport by recruiting kinesin-1 to microtubules. The Journal of biological chemistry 294, 10160–10171 (2019).

44. Hooikaas, P.J. et al. MAP7 family proteins regulate kinesin-1 recruitment and activation. J Cell Biol 218, 1298–1318 (2019).

45. Ferro, L.S. et al. Structural and functional insight into regulation of kinesin-1 by microtubule-associated protein MAP7. Science 375, 326–331 (2022).

46. Castoldi, M. & Popov, A.V. Purification of brain tubulin through two cycles of polymerization-depolymerization in a high-molarity buffer. Protein Expr Purif 32, 83–88 (2003).

47. Kellogg, E.H. et al. Insights into the Distinct Mechanisms of Action of Taxane and Non-Taxane Microtubule Stabilizers from Cryo-EM Structures. J Mol Biol 429, 633–646 (2017).

48. Widlund, P.O. et al. One-step purification of assembly-competent tubulin from diverse eukaryotic sources. Mol Biol Cell 23, 4393–4401 (2012).

49. Minoura, I. et al. Overexpression, purification, and functional analysis of recombinant human tubulin dimer. Febs Lett 587, 3450–3455 (2013).

50. Gaitanos, T.N. et al. Peloruside A does not bind to the taxoid site on β-tubulin and retains its activity in multidrug-resistant cell lines. Cancer Res 64, 5063–5067 (2004).

51. Ti, S., Ghani, V.G., Pamula, M.C. & Kapoor, T.M. Human tubulin isotypes A1B/B2B and A1B/B3 modulate sensitivity to MAP-dependent microtubule depolymerization. Mol Biol Cell 28 (2017).

52. Roostalu, J. et al. The speed of GTP hydrolysis determines GTP cap size and controls microtubule stability. Elife 9 (2020).

53. Chen, J. et al. alpha-tubulin tail modifications regulate microtubule stability through selective effector recruitment, not changes in intrinsic polymer dynamics. Developmental cell (2021).

54. Vemu, A., Garnham, C.P., Lee, D.Y. & Roll-Mecak, A. Generation of Differentially Modified Microtubules Using Enzymatic Approaches. Method Enzymol 540, 149–166 (2014).

55. Serrano, L., de la Torre, J., Maccioni, R.B. & Avila, J. Involvement of the carboxyl-terminal domain of tubulin in the regulation of its assembly. Proceedings of the National Academy of Sciences 81, 5989–5993 (1984).

56. Duan, J.M. & Gorovsky, M.A. Both carboxy-terminal tails of α- and β-tubulin are essential, but either one will suffice. Curr Biol 12, 313–316 (2002).

57. Chew, Y.M. & Cross, R.A. Taxol acts differently on different tubulin isotypes. Commun Biol 6 (2023).

58. Luo, J. et al. An evolution-conserved allosteric network in human tubulin governs paclitaxel efficacy. Nature Chemical Biology (2026).

59. Paquette, A.L. et al. Competition for microtubule lattice spacing between a microtubule expander and compactor. Curr Biol 35, 4442–4452 (2025).

60. Ettinger, A., van Haren, J., Ribeiro, S.A. & Wittmann, T. Doublecortin Is Excluded from Growing Microtubule Ends and Recognizes the GDP-Microtubule Lattice. Curr Biol 26, 1549–1555 (2016).

61. Khushalani, D.M., Kar, J., Nayak, S.B., Chaudhary, S.C. & Mohan, N. MAP4-MAP7D1 partitioning on tyrosinated-detyrosinated microtubules coordinates lysosome positioning in nutrient signalling. bioRxiv (2025).

62. Tan, R. et al. Microtubules gate tau condensation to spatially regulate microtubule functions. Nature cell biology 21, 1078–1085 (2019).

63. Beaudet, D., Berger, C.L. & Hendricks, A.G. (eLife Sciences Publications, Ltd, 2026).

64. Hoover, B.R. et al. Tau Mislocalization to Dendritic Spines Mediates Synaptic Dysfunction Independently of Neurodegeneration. Neuron 68, 1067–1081 (2010).

65. Fan, X., Okada, K., Lin, H., Ori-McKenney, K.M. & McKenney, R.J. A pathological phosphorylation pattern enhances tau cooperativity on microtubules and facilitates tau filament assembly. Res. Sq. (2025).

66. Monroy, B.Y. et al. Competition between microtubule-associated proteins directs motor transport. Nature communications 9, 1487 (2018).

67. Hotta, T. et al. (eLife Sciences Publications, Ltd, 2026).

68. Zehr, E.A., Sun, S.F., Sarbanes, S.L. & Roll-Mecak, A. Microtubules in the axon are GDP bound but adopt a stable GTP-like expanded state. Nat Struct Mol Biol (2026).

69. Yue, Y., Hotta, T., Higaki, T., Verhey, K.J. & Ohi, R. Microtubule detyrosination by VASH1/SVBP is regulated by the conformational state of tubulin in the lattice. Curr Biol 33, 4111-+ (2023).

70. Chen, J.Y. et al. α-tubulin tail modifications regulate microtubule stability through selective effector recruitment, not changes in intrinsic polymer dynamics. Dev Cell 56, 2016-+ (2021).

71. Hyman, A. et al. Preparation of Modified Tubulins. Methods in Enzymology 196, 478–485 (1991).

72. Mastronarde, D.N. SerialEM: A Program for Automated Tilt Series Acquisition on Tecnai Microscopes Using Prediction of Specimen Position. Microscopy and Microanalysis 9, 1182–1183 (2003).

73. Zhang, R. & Nogales, E. A new protocol to accurately determine microtubule lattice seam location. Journal of Structural Biology 192, 245–254 (2015).

74. Cetin, B. et al. Structure and mechanism of microtubule stabilization and motor regulation by MAP9. bioRxiv, 2025.2011.2017.688911 (2025).

75. Punjani, A., Rubinstein, J.L., Fleet, D.J. & Brubaker, M.A. cryoSPARC: algorithms for rapid unsupervised cryo-EM structure determination. Nature Methods 14, 290–296 (2017).

